# Repeated malaria exposure reprograms Vδ2+ γδ T cell chromatin accessibility and immune function toward a reversible hyporesponsive state

**DOI:** 10.64898/2026.10.02.755737

**Authors:** Kathleen D. Press, Midhuna I. Joseph Maran, Adam Ashkar-Masarwa, Kylie Camanag, Derek Chen, Vida Shokoohi, Arwa Kathiria, Maureen Ty, Jason Nideffer, John Rek, Joaniter Nankabirwa, Felistas Nankya, Julia Schulze-Hentrich, Isaac Ssewanyana, Moses Kamya, Michelle Boyle, Bryan Greenhouse, Grant Dorsey, Margaret Feeney, Will Greenleaf, Sandy Klemm, Fabian Müller, Prasanna Jagannathan

## Abstract

Vδ2+ γδ T cells contribute to antimalarial immunity via pro-inflammatory and cytotoxic mechanisms. Among children who have experienced repeated episodes of malaria, Vδ2+ T cells develop reduced pro-inflammatory function through mechanisms that remain unknown. We hypothesized that recurrent malaria drives coordinated epigenetic and transcriptional remodeling of these semi-innate cells. Using PBMC from Ugandan children in cohorts spanning high and low transmission (10 children) and longitudinally across a marked decline in transmission (20 children, 3 timepoints), we profiled the epigenome and transcriptome of Vδ2+ T cells by paired bulk ATAC-seq and RNA-seq and by CITE-seq. Higher malaria exposure was associated with reduced Vδ2+ T cell TNFα and IFNγ production and with thousands of differentially accessible chromatin regions. Immunoregulatory genes including *LYN*, *PLCG2*, and *HAVCR2* (Tim-3) showed increased chromatin accessibility and/or expression at high exposure, whereas pro-inflammatory gene (*IFNG, GZMK, IL7R*) and AP-1 (JUN/FOS) motif accessibility were increased at low exposure; linkage of regulatory elements with target genes corroborated these associations. Single-cell transcriptional analysis identified a cytotoxic CD8+CD16+CD57+ Vδ2+ T cell subset. In our *in vitro* model, repeated stimulation with malaria-infected red blood cells or the phosphoantigen HMBPP was similarly associated with reduced Vδ2+ T cell degranulation and cytokine production. Diminished responsiveness was predominantly TCR-dependent, accompanied by heightened p38 signaling, and was partially reversed by rest induced by the tyrosine kinase inhibitor dasatinib. Together these data reveal how repeated malaria may reshape cellular immunity by reprogramming Vδ2+ T cells toward a reversible, hyporesponsive state and highlight targets for therapeutically restoring their function.

## Introduction

Despite representing only a small percentage of T cells (2-5%), γδ T cells are a unique cell type that have both innate- and adaptive-like roles in protection from numerous infections. These cells uniquely recognize a broad range of antigens—including phosphoantigens in the case of the Vy9^+^Vδ2^+^ subset, the predominant population in circulation in humans—without the need for major histocompatibility complex (MHC). In the context of *Plasmodium falciparum (Pf)* malaria, which remains a major global health problem causing 600,000 deaths per year (*1*), γδ T cells expand rapidly in response to *in vitro* or *in vivo* malaria exposure (*2-4*). During primary infection, Vy9^+^Vδ2^+^ (Vδ2^+^) T cells produce pro-inflammatory cytokines and kill *Pf* parasites via a process that is granulysin dependent, and recent evidence suggests that they can also phagocytose *Pf*-infected red blood cells (*5*). Higher frequencies of Vδ2^+^T cells are associated with lower parasite densities and risk of subsequent infection in natural infection (*6, 7*), as well as protection from re-infection following vaccination with whole sporozoite vaccines (*8-11*). Understanding the contribution of Vδ2^+^ γδ T cells to antimalarial immunity, which involves a transition from symptomatic infections to predominantly asymptomatic infections, is essential to improving the efficacy of current vaccines and reducing malaria-caused mortality and morbidity.

Interestingly, children who have experienced repeated symptomatic *Pf* infections have Vδ2^+^ γδ T cells characterized by reduced expression of pro-inflammatory cytokines and proliferation in response to *Pf* parasites, as well as increased expression of immunoregulatory genes such as *FCGR3A* (CD16), *B3GAT1* (CD57), *HAVCR2* (Tim-3), *BATF, LYN, PLCG2,* and KIRs (*6, 7, 12, 13*). In contrast, Vδ1+ γδ T cells, which are more common in mucosal tissue and lymphoid organs, continue to expand and become cytotoxic and activated following repeated infections (*14, 15*). Activated Vδ2^+^T cells from individuals with chronic malaria exposure respond to opsonized antigen via CD16 in a pathway independent of the T cell receptor (TCR) (*16*). Furthermore, although higher pro-inflammatory Vδ2^+^ γδ T cells responses have been associated with protection against infection (*7*), loss of this pro-inflammatory response has been associated with a greater ability to tolerate malaria parasites without symptoms (*6*). Although mechanisms driving this altered γδ T cell function have remained unknown, several innate immune cells have been shown to develop a form of immune “memory” following antigenic exposure, driven in part by epigenetic and transcriptional changes (*17, 18*). In monocytes, this includes both “trained immunity” (increased responsiveness to malaria)(*19, 20*) and tolerance (decreased responsiveness)(*21, 22*), including histone methylation at immunometabolic promoters associating with altered responsiveness to toll like receptor stimulation (*20*). Genomic sites of chromatin accessibility—which comprise candidates for transcription factor binding and gene transcription regulation—can be assessed using ATAC-seq, a method using Tn5 transposase to insert sequencing adapters at sites of open chromatin (*23*). Whether repeated malaria exposure conditions epigenetic and transcriptional remodeling of semi-innate γδ T cells remains unclear.

In this study, we utilized samples from children enrolled in regional and longitudinal cohorts in Uganda, as well as novel in *in vitro* stimulation assays to characterize the function, epigenome, and transcriptome of Vδ2^+^ T cells following repeated malaria exposure. We hypothesized that associations between increased malaria incidence and altered Vδ2^+^ T cell function (*6, 7*) are driven by transcriptional and epigenetic reprogramming. Improving our understanding of the inflammatory and immunoregulatory roles of Vδ2^+^ T cells during malaria infection may provide opportunities to manipulate this response therapeutically.

## Results

### Malaria exposure correlates with altered Vδ2^+^ T cell chromatin accessibility at genomic sites associated with the regulation of inflammation and exhaustion

Building on prior studies showing that repeated malaria is associated with reduced function of γδ T cells, we re-analyzed published data on Ugandan Vδ2^+^ γδ T cell responses (*7*), comparing cytokine production among Vδ2^+^ T cells from children living in two different malaria transmission settings **(Fig 1A, Table S1)**. Differential transmission was assessed using household-level entomological inoculation rate (aEIR), a calculation estimating annual exposure to infected mosquitoes using data from mosquito captures in the homes of each cohort participant (*24*). Peripheral blood mononuclear cells (PBMC) from children in Tororo (aEIR=310 in 2014) and Jinja (aEIR=2.8 in 2014) were stimulated for 24 hours with *Pf*-infected red blood cells (iRBC). Vδ2^+^ T cells pro-inflammatory cytokine (TNFα, IFNγ) production in response to stimulation with iRBC was significantly lower among children living in Tororo compared to Jinja **(Fig 1B-C)** and significantly correlated with aEIR **(Fig S1A)**.

**Figure 1:**
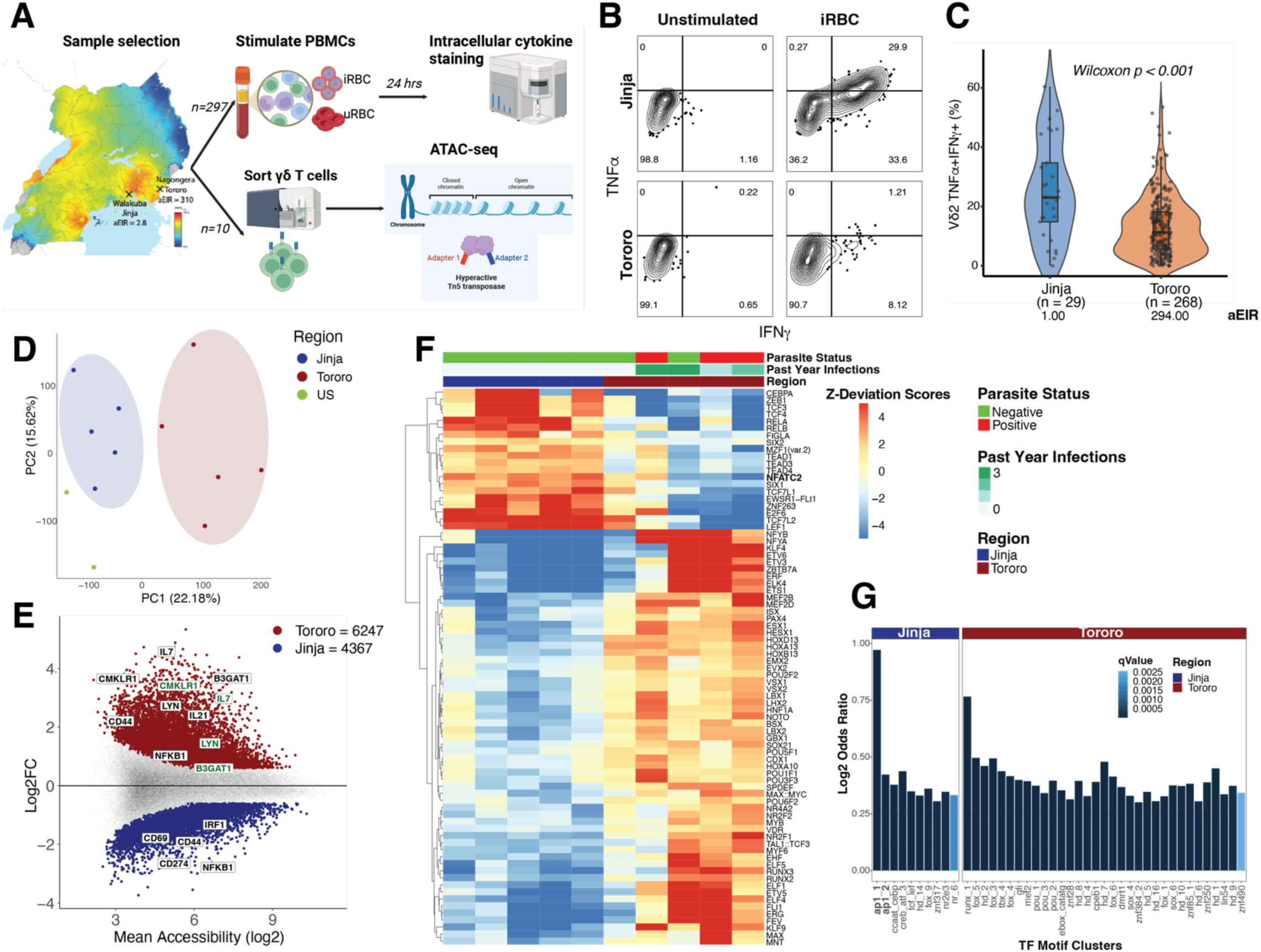
Malaria exposure correlates with altered Vδ2+ T cell cytokine production and chromatin accessibility at sites associated with the regulation of inflammation and exhaustion. **A.** Schematic of study design. Created in https://BioRender.com **B-C.** PBMC from children living in Tororo (highly endemic region) or Jinja (low transmission) were stimulated with iRBC for 24 hours. (B) Representative flow cytometry plot. (C) Production of TNFα and IFNγ by Vδ2^+^ T cells in response to iRBC were significantly lower among children living in Tororo compared to children living in Jinja. **D.** PCA plot showing separation of samples by region. Variance explained by PC1 and 2 is annotated in the axis labels. **E.** MA plot of differentially accessible peaks (DESeq2) between regions Jinja and Tororo (padj 0.05, logFC 0.6, N=5 for each group). Green indicates proximal gene (<=2k from TSS) and black distal gene (2k > distal > 250k). **F.** Heatmap of ChromVAR’s significantly differential motifs between Jinja and Tororo (Limma eBayes, padj < 0.05, |log2FC|> 0.6). **G.** Bar plots showing motif families (*26*) that are enriched by LOLA analysis (Fisher’s Exact Test, qval < 0.05, logOR > 0.3).

To test whether functional changes in Vδ2^+^ T cells after repeated malaria antigen exposure associated with epigenetic reprogramming, we performed ATAC-seq with sorted Vδ2+ and CD4+ T cells from children of the same age group in Tororo compared to Jinja, as well as US controls **(Fig 1A**, **Fig S1B-C)**. Variation in Vδ2+ T cell chromatin accessibility was largely explained by transmission region, and Jinja samples were more similar to US controls than Tororo samples **(Fig 1D)**. We identified 10,614 differentially accessible regions (DARs) between Vδ2^+^ T cells from children in Tororo *vs.* Jinja **(Fig 1E**, **Table S2)**, with the majority hyperaccessible in the Tororo (high transmission) samples (6247 vs. 4367 DARs). DARs represent genomic regions potentially acting as candidate cis-regulatory elements (cCREs) for their target genes. Associating the 10,614 DARs to their 5176 closest genes also indicated more accessible regulators of genes in the Tororo samples (3591 vs. 2218 nearest genes). Overall, DARs that were more accessible in Tororo (high exposure) were more indicative of an exhausted-like state (*B3GAT1*) or alternate inflammatory pathways (*IL17, LYN, CMKLR1, IL21*) while those that were more accessible in Jinja (low exposure) were more pro-inflammatory (*NFKB, IRF1, CD44, CD69*). Notably, there was significant overlap (P = 0.00005; 31 genes) between DARs that were identified as more accessible in Tororo (high transmission) and genes that were more highly expressed among children with higher malaria incidence in our previously published microarray data (*6*) (**Table S2**) (the corresponding comparison between Jinja (low transmission) and genes associated with higher malaria incidence in the microarray was not significant (P = 0.9957; 5 genes)). Parasite status and infection within the previous year additionally associated with differential accessibility **(Fig S1D)**. Only 10 DARs were identified for CD4+ T cells **(Fig S1E)**, highlighting that the observed differential chromatin accessibility among Vδ2+ T cells was likely cell specific. When we investigated the activity of transcription factors (TFs) as quantified by the aggregate accessibility in candidate binding loci, we observed association of specific TFs and TF families (clusters of TFs with similar binding motifs) with variation in chromatin accessibility **(Fig 1F-G, Fig S1F).** For example, NFATC2 **(Fig 1F)** and AP-1 family binding sites **(Fig 1G)** –both of which are important modulators of inflammation—were highly accessible in the Jinja samples. Footprinting analysis further highlighted differential transcription factor binding profiles, including FLI1, whose motifs were more accessible in Tororo (high transmission) samples **(Fig S1G)**, and which has shown association with CD8+ T cell exhaustion (*25*). Associations between malaria exposure and differentially accessible chromatin proximal to Vδ2+ T cell genes involved in immune regulation (e.g. *B3GAT1, IL17* and *LYN* more accessible at high exposure) and inflammation (e.g. *NFKB, IRF1*, and AP-1 family more accessible at low exposure) supports the hypothesis that recurrent malaria may impact Vδ2+ T cell functional activity via epigenetic modulation.

### High malaria transmission in longitudinal samples associates with increased Vδ2+ T cell chromatin accessibility at immunoregulatory genes while low transmission associates with increased accessibility at inflammatory genes and AP-1 factor motifs

Given that our cross-sectional data strongly supported a gene-regulatory role of chromatin accessibility in immune responsiveness, we next sought to determine whether Vδ2^+^ T cell functional, epigenetic and transcriptional changes could be observed within individuals over time. While historically malaria transmission in Tororo was among the highest in the world, transmission dramatically decreased towards the end of 2014 following district-wide insecticide spraying campaigns leading to reduced parasite prevalence and disease incidence **(Fig 2A)**(*27*). Vδ2^+^ T cells from the same Tororo children exhibited higher responsiveness to 6-hour stimulation with iRBCs in 2015 (when transmission was low after insecticide spraying) compared to 2013 (when transmission was high before insecticide spraying), as indicated by higher degranulation capacity (CD107a) and production of TNFα and IFNγ **(Fig 2B)**.

**Figure 2:**
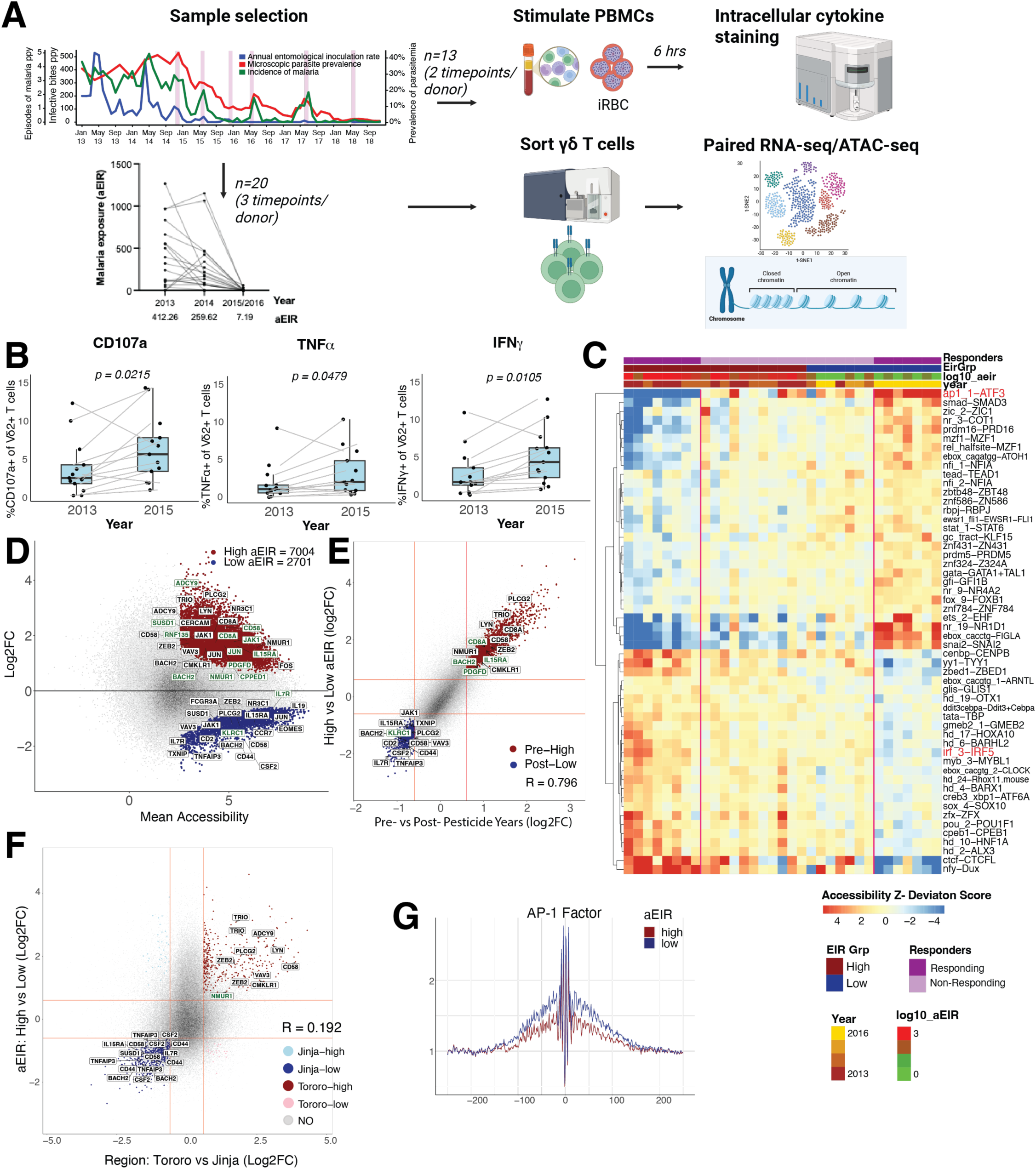
High malaria transmission associates with increased Vδ2+ T cell chromatin accessibility at immunoregulatory genes while low transmission associates with increased accessibility at inflammatory genes and AP-1 factor motifs. **A.** Schematic showing experimental design for stimulation and paired RNA-seq/ATAC-seq experiments. Created in https://BioRender.com Top left panel (adapted from (*30*)): Malaria incidence and exposure in Tororo, Uganda changed dramatically between 2013 – 2018. Pink bars indicate insecticide spraying campaigns. **B.** PBMCs from the same children (n=13) in 2013 and 2015 were stimulated for 6 hours with malaria-infected red blood cells (iRBCs). Vδ2^+^ T cells at the 2015 timepoint expressed significantly more CD107a, TNFα a and IFNγ than at the 2013 timepoint. **C.** Heatmap of ChromVAR’s significantly differential (donor corrected, Wilcoxon Test, padj < 0.05, |delta mean| > 1) motif clusters using intra-aEIR group k-means clustering (n=33; donors with at least sample in both low and high aEIR groups were included). Red lines demarcate responder from non-responder samples. **D.** MA plot of differentially accessible peaks (Paired DESeq2 analysis) between high and Low aEIR of responder samples (padj < 0.05, |log2FC| > 0.6). High aEIR corresponds to >100. Green indicates proximal gene (<=2k from TSS) and black distal gene (2k > distal > 250k). **E.** Scatterplot of differentials showing high correlation between aEIR (low vs high; responders only) and insecticide year-based (pre vs post insecticide; all paired samples) analyses. Green indicates proximal gene (<=2k from TSS) and black distal gene (2k > distal > 250k). **F.** Scatterplot showing correlation between differential accessibility between longitudinal samples (low vs high aEIR) and regional data (Tororo vs Jinja) analyses. Gene names indicate closest gene by distance. **G.** Transcription Factor Footprint plot of AP-1 factor motif family, showing more accessibility in low aEIR around TF binding sites.

To better understand the gene-regulatory mechanisms driving these changes on a genome-wide scale, we performed bulk paired RNA-seq and ATAC-seq experiments among Vδ2+ T cells from 20 Ugandan children at 3 timepoints – 2 prior to insecticide spraying (2013 and 2014) and one after (2015 or 2016) **(Fig 2A**, **Fig S2A-D**, **Table S3)**. By using multiple samples from each child, including before and after the decline in malaria transmission, we could assess the relationship between chromatin accessibility, transcription, and household-level exposure to *Pf*-infected mosquitoes (aEIR) (**Fig S2E**). Analyses of clinical or exposure-related differential accessibility by ATAC-seq (including pairwise group comparisons based on year, aEIR, malaria incidence, and parasite status) were performed across subsets of paired samples among the 46 samples retained following QC **(Fig S3A-B)**. We identified 3,936 candidate cis-regulatory elements (cCREs) that were significantly differentially accessible between timepoints when regional malaria transmission was high (2013/2014) compared to when transmission was low (2015/2016) (**Fig S3A, Table S4**, N=40). Accessibility at 770 cCREs (corresponding to 656 nearest genes) associated with higher household-level aEIR while 185 DARs (173 genes) associated with low aEIR **(Fig S3A-B, Table S4**, N=33). While aEIR and year are closely related **(Fig S2E)**, we focused on aEIR for the remaining analysis given that aEIR is a more precise measure of transmission and is not confounded by age.

We characterized the chromatin accessibility dynamics in transcription factor family binding motifs across the 33 samples that included matching samples from each donor across exposure groups **(Fig 2C)**. Intra-aEIR group k-means clustering of the differentially accessible motifs analysis revealed 3 visually identifiable clusters: 1) samples from high EIR timepoints with clear increased accessibility at binding sites for a specific set of TF motifs, 2) samples without strong differential TF activity, and 3) samples from low EIR timepoints with increased TF activity for a specific set of motifs **(Fig 2C)**. Differences in aEIR between high and low timepoints were larger for the 14 samples in clusters 1 and 3 (“responders”) compared to the samples in cluster 2 (“non-responders”) **(Fig S4A)**. Increased aEIR associated with reduced chromatin accessibility at AP-1 factor (e.g. JUN, FOS families) binding sites **(Fig 2C, Fig S4B-C)** and increased accessibility at sites associated with the IRF3 motif family (which contains 3 motif models for IRF5, a regulator of the type I interferon inflammatory pathway) **(Fig 2C, Fig S4C)**; the former matches results from the regional study. Given that comparing groups 1 and 3 captured the strongest differential signature in agreement with other quantifications of exposure **(Fig S3B)**, we focused on these “responders” (13 individuals with at least one sample per exposure group) for the remainder of the analysis.

Among this responder group, we identified 7,004 DARs (corresponding to 5,479 nearest genes) whose accessibility associated with higher household-level aEIR and 2,701 (1,584 genes) associated with lower aEIR **(Fig 2D, Table S4)**. These DARs overlapped significantly with those identified in the year-wise comparison (2013/2014 (pre) vs. 2015/2016 (post)) **(Fig 2E)**, which is consistent with the high association between year and aEIR **(Fig S2E)**. Numerous genes of interest that are involved with immune signaling or regulation were linked to DARs more highly accessible at high aEIR timepoints, including *CD8, IL19, TNFAIP3, CMKLR1, LYN, ZEB2, HAVCR1*, and *BATF*. Nine genes, including *CD8, CMKLR1, LYN*, and *BATF* overlapped with those previously seen as upregulated in children with higher malaria exposure (*6*) and CD8 was also recently described as a marker of effector γδ T cells in the context of chronic tuberculosis infection (*28*). Nearest genes to DARs that were more accessible for low aEIR included *CSF2, BACH2, IL15RA, NR3C1, JAK1, CD44, KLRC1, IL7R, CD2, CD58*, and *CCR7*. Together, these point to a more hyporesponsive state when transmission is high and a pro-inflammatory phenotype that is regained when transmission decreases.

Comparing the cross-sectional study with the longitudinal, overlap between DARs from Jinja (low transmission) and low aEIR in the longitudinal study was statistically significant (p<0.00005, hypergeometric test; overlap for Tororo/high aEIR not significant). Overlapping DARs included *CD44, IL7R, IFNG*, and *GZMK* for Jinja and longitudinal low aEIR, *and LYN, PLCG2, and CMKLR1* for Tororo and longitudinal high aEIR **(Fig 2F)**. The increased accessibility at IFNG for both Jinja and low EIR is consistent with increased TNFα and IFNγ production from samples at a low transmission timepoint compared to high transmission (**Fig 2B**). Additionally, there was overlap in differential transcription factor activity between the cross-sectional and longitudinal studies **(Fig S4D-E)**. Footprinting analysis confirmed differential accessibility of motifs for transcription factors associating with malaria exposure, including the AP-1 factor family **(Fig 2G, Fig S4F**), which was more accessible among low EIR samples. These results suggest that similarly to how c-Jun deficiency has been linked to T cell exhaustion while c-Jun overexpression attenuates exhaustion (*29*), AP-1 immune pathways could be modulated by repeated malaria exposure to cause downstream hyporesponsiveness in Vδ2+ T cells.

### Decreased malaria exposure in longitudinal samples is associated with transcriptional shifts towards a restored pro-inflammatory phenotype in Vδ2+ T cells

Paired RNA-seq of the sample set from pre- and post-insecticide timepoints in the longitudinal cohort **(Fig S5A)** identified 149 differentially expressed genes (DEGs) between high and low aEIR (70 more highly expressed in high aEIR samples and 79 more highly expressed in the low aEIR group) **(Fig 3A-B**, **Table S5)**. Genes upregulated in the high aEIR group included immune regulatory genes such as *HAVCR2, KIR3DL2, KIR2DL4, PLCG2, CD8A, KIR2DL1, LYN, KIR3DL1, IL10,* and *CD244*, while genes upregulated at low aEIR samples were enriched in markers of inflammation and killing (i.e. *CD44, TNF, IFNG, IL7R, GZMK*, and *IL23R*). Seven genes, including *HAVCR2, LYN, PLCG2*, and *CD8* overlapped with those previously seen as upregulated in children with more malaria infections (*6*) and the upregulation of *CD8* and KIR receptors was also consistent with observations of γδ T cells following chronic tuberculosis infection (*28*). Differential gene expression and differential gene activity defined by the aggregate accessibility of genes and associated accessible regions exhibited low-moderate correlation (including concurrences of genes such as *CD44, IL7R* and *IFNG*) **(Fig 3C),** indicating that their expression is likely modulated by epigenetic mechanisms. Numerous pathways associated with positive regulation and activation of immune responses (e.g. regulation of immune process, lymphocyte activation, cell killing, cell adhesion, lymphocyte-mediated immunity, cytokine production) were enriched in genes with increased expression in the low aEIR samples while pathways associated with negative immune regulation (e.g. inhibition of signal transduction and cell surface receptor signaling pathways, negative regulation of lymphocyte proliferation) were enriched in genes upregulated in the high aEIR group **(Fig S5B-C)**. Comparison of the differentially expressed (DEGs) with transcription factor binding motif activity (ChromVAR) revealed concurrent increases in accessibility and expression between the high and low aEIR groups **(Fig S5D).** DEGs were also identified based on year, malaria incidence, and parasite status (**Fig S6A-B**), suggesting that parasite exposure and symptomatic infection interact to impact gene regulation. Together, differential gene expression and chromatin accessibility between samples representing high and low malaria exposure levels support a role for coordinated transcriptional and epigenetic regulation of numerous immune regulation pathways in modulating differential Vδ2+ T cell function.

**Figure 3:**
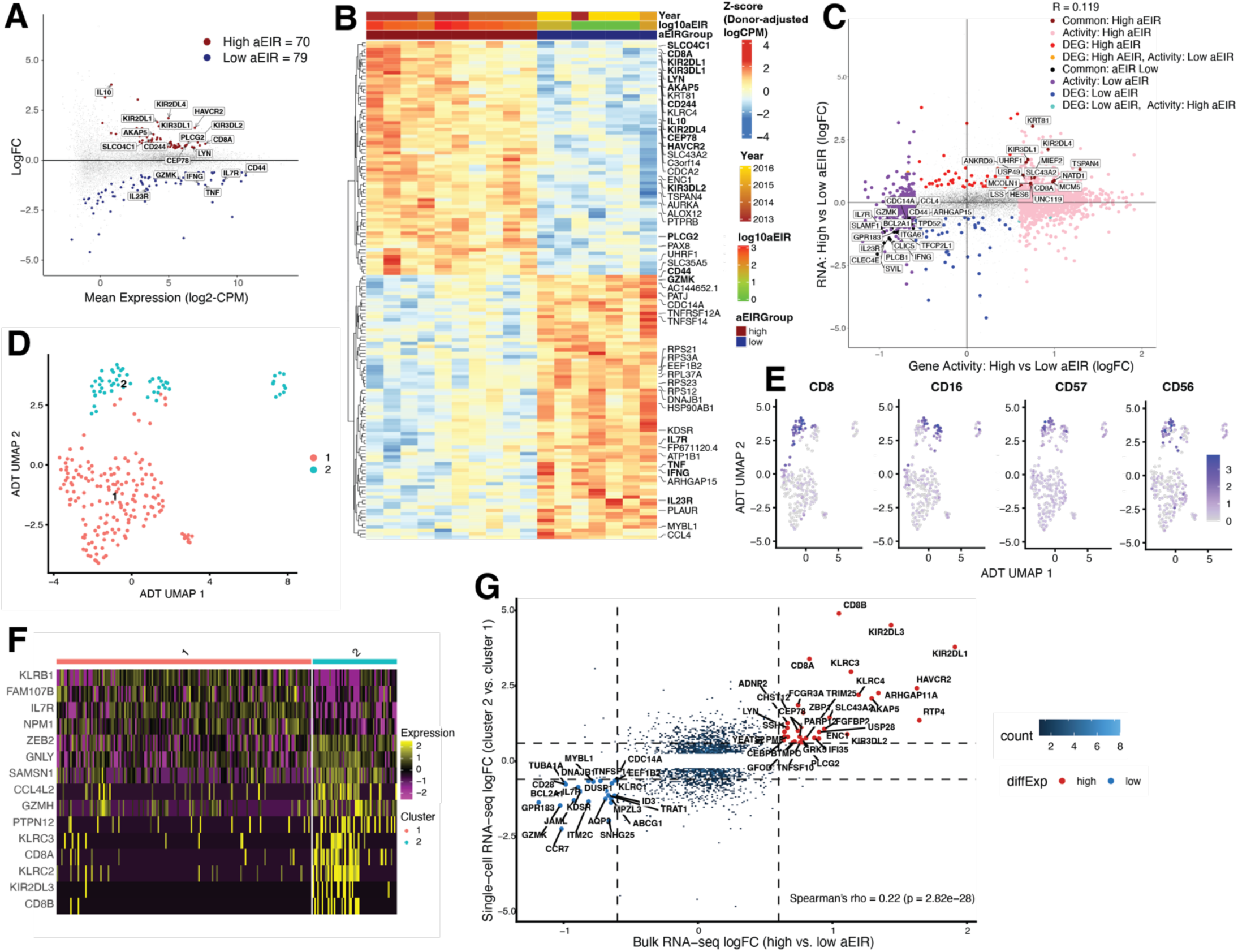
High malaria transmission and a CD8+CD16+CD57+ cell population both associate with increased Vδ2+ T cell expression of immunoregulatory genes like *HAVCR2, LYN* and *PLCG2* and reduced expression of *IL7R*. **A.** MA plot of differential expression analysis comparing high (>100) vs low (<100) inoculation rate (aEIR). The differentially expressed genes (DEGs) are colored in red (high aEIR) and blue (low aEIR) (logFC 0.6, padj 0.05, EdgeR’s Likelihood Ratio Test (LRT). N=10 for High and 7 for Low. **B.** Heatmap of the DEGs from (A). **C.** Agreement of differentially regulated genes identified by the gene activity (calculated from ATAC data) and gene expression from (A, B). **D.** ADT-based UMAP projection of Vδ2 T cells based colored by ADT-based Louvain clusters **E.** Feature plots showing normalized ADT expression of selected surface markers. **F.** Heatmap showing the top differentially expressed genes between Vδ2+ T cell ADT-based cluster 1 and 2 **G.** Scatter plot showing log_2_ fold changes from comparing differential expression of high *vs.* low aEIR groups in the bulk analysis (x-axis) and cluster 2 *vs.* cluster 1 in the CITE-seq analysis (y-axis). Each point represents a gene. Red and blue points indicate genes significantly upregulated or downregulated in the bulk high aEIR group, respectively. Dashed lines indicate the significance and fold-change thresholds used for differential expression. Genes in the upper right quadrant are enriched in high aEIR cells and cluster 2, whereas genes in the lower left quadrant are enriched in low aEIR cells and cluster 1. P-value calculated using Fisher’s Exact Test.

### Single-cell transcriptional analysis identifies a cytotoxic CD8+CD16+CD57+ Vδ2+ T cell subset

To determine whether specific subsets of Vδ2+ T cells drive the differential transcription observed in bulk analysis, we used CITE-seq, which enables simultaneous detection of surface protein abundance and gene expression in single cells. Our analysis included samples from 3 Ugandan children in Tororo at 2 timepoints (2013 and 2015) **(Table S6)**. After applying global filtering thresholds based on RNA and protein characteristics (**Fig S7A**) and batch correction (**Fig S7B**), cell types were identified by Louvain clustering (**Fig S7C**). Paired ADT measurements were used to identify the subset of cells expressing the Vδ2+ TCR marker (**Fig S8A-E**). Unsupervised analysis of these Vδ2+ T cells identified two phenotypically distinct clusters (**Fig 3D**). Cluster 2 exhibited elevated expression of CD8, CD16, CD57 and CD56 **(Fig 3E, Fig S9A**), consistent with a phenotype of chronic antigen exposure (*28*). Differential expression analysis showed that compared to all other Vδ2+ T cells, Cluster 2 was distinguished transcriptionally by decreased expression of *KLRB1*, *IL7R* and *GZMK* and increased expression of *GNLY, GZMH, KIR2DL3*, and *ZEB2* (**Fig 3F, Fig S9B**). Gene Ontology (GO) enrichment analysis revealed that genes upregulated in cluster 2 associate with cytotoxicity while downregulated genes associate with T-cell differentiation and migration **(Fig S9C-D)**. A subpopulation of CD8+ γδ T cells in the setting of chronic tuberculosis similarly upregulated *CD16, GNLY, GZMH*, and several KIRs despite reduced TCR responsiveness (*28*) and IFNγ-producing CD16+ γδ T cells also increase in response to cytomegalovirus (*31*). When we compared the genes differentially expressed between clusters 1 and 2 with the genes that were differentially expressed between low *vs.* high aEIR groups in our bulk analysis, we observed significant overlap (Spearman’s rho = 0.22, p = 2.82e-28), including upregulation of *CD8, HAVCR2, LYN, PLCG2* and several KIRs for cluster 2 and high aEIR and upregulation of *IL7R* and *GZMK* for cluster 1 and low aEIR **(Fig 3G)**. This overlap suggests that cluster 2 could represent a phenotype associated with higher malaria exposure that exhibits non-TCR mediated cytotoxicity.

### Peak-to-gene linkage analysis identifies immunoregulatory sites in malaria exposure where accessibility correlates with target expression, including IL7R linked with low exposure

In order to identify cCREs linked to differentially regulated genes, we next conducted a peak-to-gene analysis between differentially expressed genes (DEGs) and differentially accessible regions (DARs) by aEIR group (high vs low) **(Fig 4A)**. We identified 132 genes for which accessibility at gene-proximal and gene-distal cCREs and gene expression were correlated (**Fig 4B, Table S7)**. Particular genes of interest associated with high exposure included genes that downregulate inflammatory responses (*IL10*), were previously shown to be upregulated in children with more malaria infections (*LYN, PLCG2, CD8A, HAVCR2*) (*6*), or play a role in NK cell activation/inhibition (*NCR1, KLRC4, KIR2DL1, KIR2DL4, KIR3DL1, KIR3DL2*) **(Fig 2B)**. Genes of interest associated with low exposure included those that associate with activation (*CD44*) or upregulate inflammation and cytotoxicity (*IL7R, TNF, IFNG, GZMK*) **(Fig 2B)**; association of *TNF* and *IFNG* with low exposure additionally supports our observation that samples from children with lower malaria exposure produce more TNFα a and IFNγupon secondary stimulation *in vitro* **(Fig1B-C, Fig 2B)**. Similarly to our observed association between KIR receptors and higher malaria exposure, previous work has shown that Vδ2+ T cells differentially express KIR inhibitory receptors based on malaria exposure and that expression of specific KIRs associate with reduced (e.g. KIR2DL1) or increased (e.g. KIR3DL1) degranulation and cytokine production (*13*). Accessibility profiles at and around the gene loci of *LYN* (associated with high aEIR) and *IL7R* (associated with low aEIR) (**Fig 4C**, **FigS10-S11)** and other loci representing these categories (**Figs S12-S14**) visually confirmed these differences. Transcription factor binding sites enriched in the linked peaks included AP-1 factors associating with low aEIR **(Fig 14C)**. Since *IL7R* was identified in the peak-to-gene analysis as well as the single-cell analysis and is known to play a role in increasing IL-17 expression (*32*), as well as in T cell effector memory response formation (*33*), we examined IL7R protein expression among 37 individuals in the longitudinal cohort. IL7R expression was significantly lower on γδ T cells collected at timepoints with high EIR compared to low-EIR timepoints **(Fig 4E)**, which supports a role for IL7R expression in Vδ2+ T cell antimalarial responses –potentially via downregulating IL-17 production and effector memory responses (*32, 33*).

**Figure 4:**
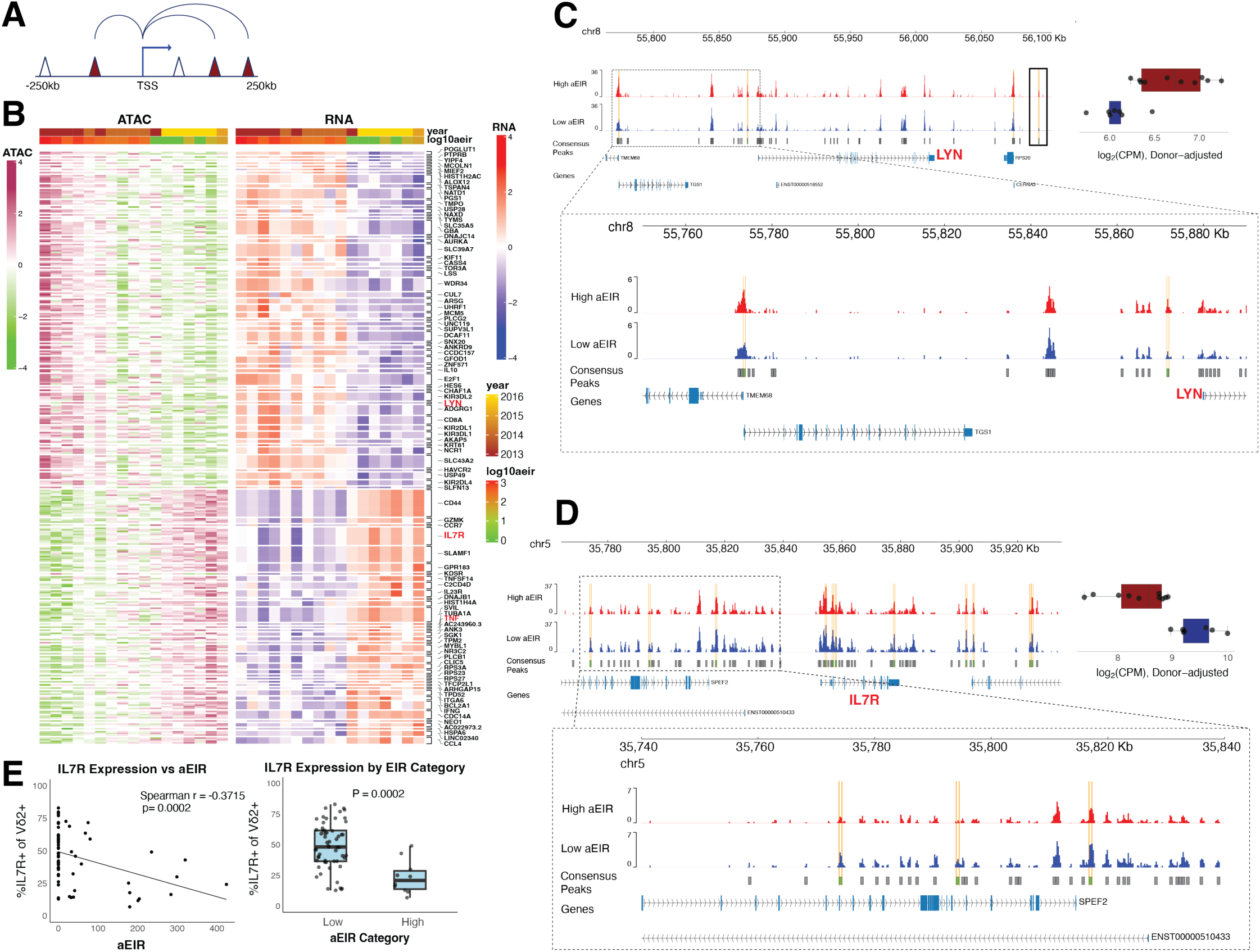
Peak-to-gene linkage analysis highlights gene expression and chromatin accessibility dependence at the TNF, IL7R, and LYN gene loci and linked cCREs. **A.** Diagrammatic representation of the Peaks to Gene correlation. The Differentially Expressed genes (DEGs) were correlated with the Differentially Accessible candidate peaks (DARs) (peaks in Red) obtained from the aEIR’s high vs low analysis. Only DARs located within 250 kb up- or downstream of a DEG’s transcription start site (TSS) were included in the analysis. **B.** Heatmap of correlated DEGs and DARs (R > 0.3). **C.** Gene track showing differential accessibility and expression for LYN between high and low EIR samples. **D.** Gene track showing differential accessibility and expression IL7R between high and low EIR samples. **E.** IL7R expression by CYTOF associates with EIR. N=70 samples from 37 individuals. High aEIR >100.

### Reduced Vδ2+ T cell degranulation and cytokine production following repeated malaria exposure is attributed to predominantly TCR-dependent mechanisms

Together, our epigenetic and transcriptional analyses identified numerous potential genes and pathways linked to altered Vδ2+ T cell function following changing malaria exposure. To better define these pathways, we established an *in vitro* assay to recapitulate elements of the *in vivo* phenotype of repeated malaria **(Fig 5A**, **Fig S15A)**. We stimulated malaria-naive PBMC or purified γδ T cells with iRBC (3:1) or the phosphoantigen HMBPP and then re-stimulated these cells 6 days later with iRBC (1:1) or positive controls. We assessed degranulation (CD107a) and pro-inflammatory cytokine production (TNFα, IFNγ). Vδ2+ T cells (whether within PBMC or purified) degranulate less and produce less cytokines in response to secondary stimulation with iRBCs after 1) stimulation with HMBPP compared to no stimulation or 2) after stimulation with iRBCs compared to uninfected red blood cells or media controls **(Fig 5B-C, Fig S15B-C)**. In contrast, 6-day stimulation with iRBCs or HMBPP does not impact the ability of the cells to respond to PMA/ionomycin **(Fig 5B-C, Fig S15B-C)**, indicating that the reduced responsiveness requires activation of the T cell receptor. Similar results for purified γδ T cells and Vδ2+ T cells within PBMC indicate that the reduced responsiveness is cell-intrinsic and not dependent on signals from other cells. When we compared RNA expression changes within samples stimulated *in vitro vs.* within the previously described longitudinal samples from Ugandan children, we identified significant overlap between genes upregulated for high aEIR and *in vitro* stimulation (e.g. *HAVCR2, KIR2DL4, KIR3DL2, LYN*) or low aEIR and lack of stimulation (*TNFRSF10A, TNFSF14, CCR7, IL7R*) **(Fig S15D)**. We repeated the stimulation assay using cells from malaria-uninfected individuals living in a malaria-endemic region in Eastern Uganda. As in the malaria-naïve samples, CD107a and TNFα responses to secondary stimulation with iRBC were significantly reduced following 6-day stimulation with iRBC or HMBPP compared to unstimulated controls **(Fig 5D).**

**Figure 5:**
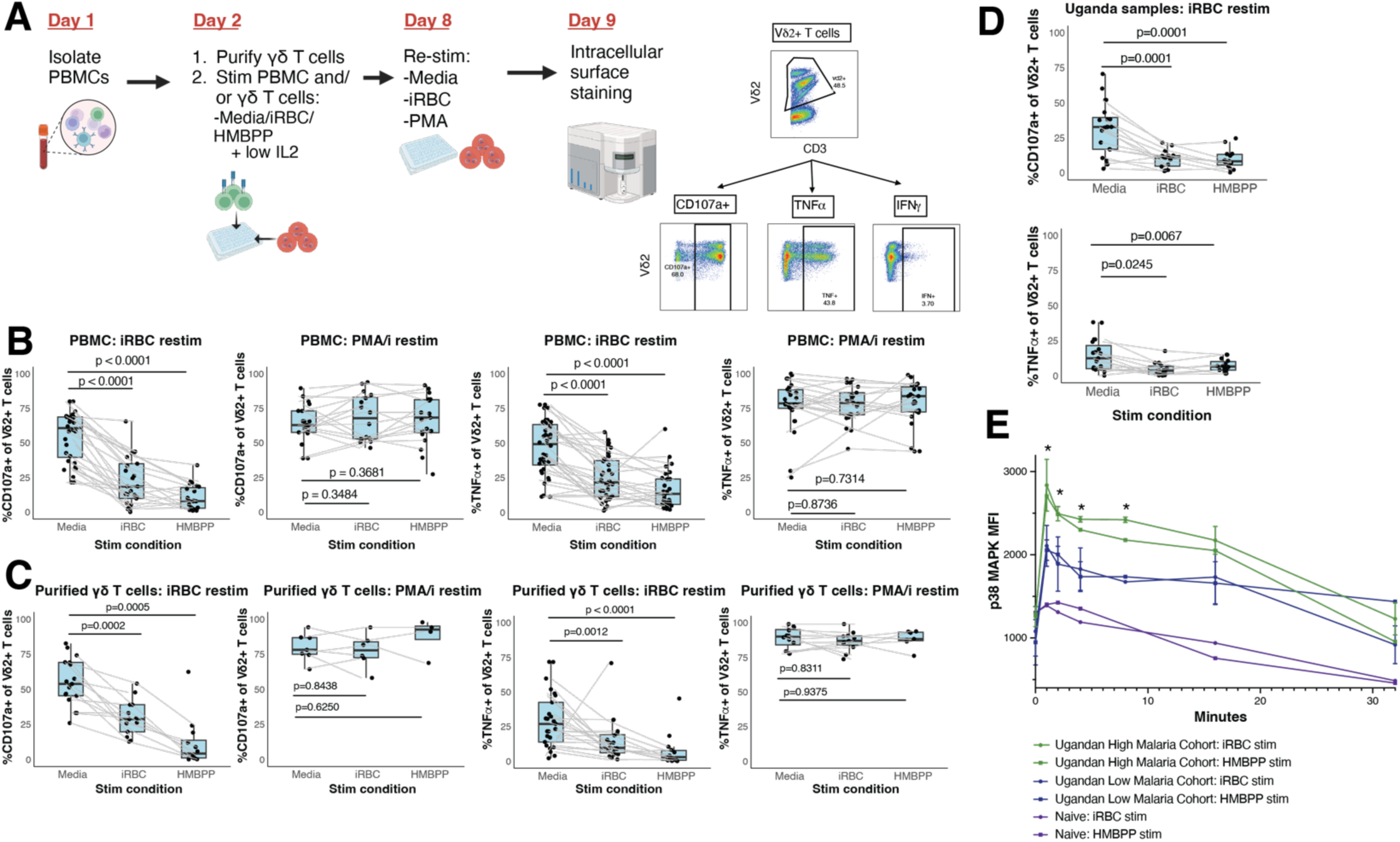
Diminished Vδ2+ T cell function following repeated malaria exposure is attributed to TCR-dependent mechanisms. **A.** Schematic showing experimental design for *in vitro* stimulation experiments. Created in https://BioRender.com **B.** Among Vδ2+ T cells within PBMCs, CD107a and TNFα production decrease following 6-day iRBC (3:1) or HMBPP stimulation followed by iRBC re-stimulation (1:1) but do not change following re-stimulation with PMA/Ionomycin. Values normalized to media re-stim condition. **C.** Among purified Vδ2+ T cells cultured with IL-2, CD107a and TNFα production decrease following 6-day iRBC or HMBPP stimulation followed by iRBC re-stimulation but do not change following re-stimulation with PMA/Ionomycin. Values normalized to media re-stim condition. **D.** PBMC from Ugandan children (n=20) were stimulated with iRBC or HMBPP for 6-days and re-stimulated with iRBC. CD107a and TNFα decrease following stimulation. Values normalized to media re-stim condition. **E.** Phosphoflow experiment showing Vδ2+ T cell p38 signaling in Uganda and naïve samples with diverse malaria exposure that were re-stimulated with iRBC.

In order to define the pathways involved in TCR-dependent activation and building off of work showing that HMBPP results in NF-κB and AP-1 activation (as well as the release of proinflammatory MIP-1α, IFNγ, and TNFα) (*34, 35*), we utilized Phosphoflow, an assay that detects phosphorylation of protein markers governing signaling pathways. When we stimulated Ugandan samples from individuals living in higher or lower malaria transmission settings with iRBCs or HMBPP, we observed that repeated malaria exposure associated with significantly higher Vδ2+ T cell p38 **(Fig 5E)**, but not pERK1/2 **(Fig S15E)**, signaling. These results suggest that repeated malaria exposure primes Vδ2+ T cells to activate p38 signaling in response to secondary stimulation. Interestingly, LYN and PLCγ2, which were both more accessible and upregulated in samples with higher malaria exposure, can be a part of p38 activation: LYN can contribute to PLCγ2 phosphorylation (*36*) and PLCγ2 can activate p38 (*37*).

The *in vitro* data above suggest that TCR-intrinsic signaling mechanisms may be driving the phenotype of reduced Vδ2+ T cell responsiveness to secondary stimulation. To further investigate the impact of repeated *in vitro* stimulation on Vδ2+ T cells, we assessed both their cell surface phenotype and whether these cells acquired other cytotoxic capabilities. Expression of the activation marker HLA-DR, as well as frequencies of effector memory cells, increased following stimulation **(Fig S16A-B)**, indicating activation and differentiation. We next tested the capacity of these samples to perform antibody-dependent cellular cytotoxicity (ADCC), which depends on expression of CD16. In contrast to increased CD16-mediated killing on CD8+ γδ T cells responding to tuberculosis (*28*), we did not find evidence that Vδ2+ T cells acquire antibody-dependent cytotoxic capacity in these short-term *in vitro* cultures **(Fig S16C)**.

### Vδ2+ T cell degranulation and cytokine production can be regained after a period of rest

Our preceding analyses showed that Vδ2+ T cells, which contribute to the control of parasite replication, become functionally hyporesponsive after repeated malaria through coordinated epigenetic and transcriptional reprogramming, and that this reduced responsiveness is predominantly TCR-dependent. Notably, Vδ2+ T cell function was greater at low-transmission timepoints, suggesting that this state may not be fixed; we therefore tested whether the hyporesponsive phenotype is reversible, a question directly relevant to the immunological consequences of declining transmission and malaria-control programs. We first exposed PBMCs to 6-day stimulation with iRBC or HMBPP followed by a 3–5-day rest period. Transient temporal rest could partially rescue Vδ2+ T cell decreased degranulation and cytokine production following iRBC re-stimulation, although this was not seen in all donors –potentially due to the rest period not being long enough **(Fig S17A)**.

It has recently been shown that functional rest of the TCR receptor can be induced with the tyrosine kinase inhibitor dasatinib, which reversibly inhibits T cell receptor signaling and effector functions (*38*) and that has been used to induce rest in exhausted chimeric antigen receptor (CAR) – T cells being developed as cancer therapeutics (*39*). We tested whether dasatinib could similarly improve immune responses in Vδ2+ T cells that have impaired function following malaria exposure **(Fig 6A)**. Dasatinib combined with HMBPP increased responses compared to HMBPP alone, revealing a reversal of the exhausted-like phenotype **(Fig 6B)**. Although dasatinib combined with iRBC did not show improved responses compared to iRBC alone **(Fig S17B)**, this may because other immune cells also take up the parasites or because tyrosine kinase inhibitors may have direct impacts on the parasite(*40*). Improved responses when dasatinib was combined with HMBPP indicates that reduced Vδ2+ T cell responsiveness could potentially be modulated by drugs to restore immune function.

**Figure 6:**
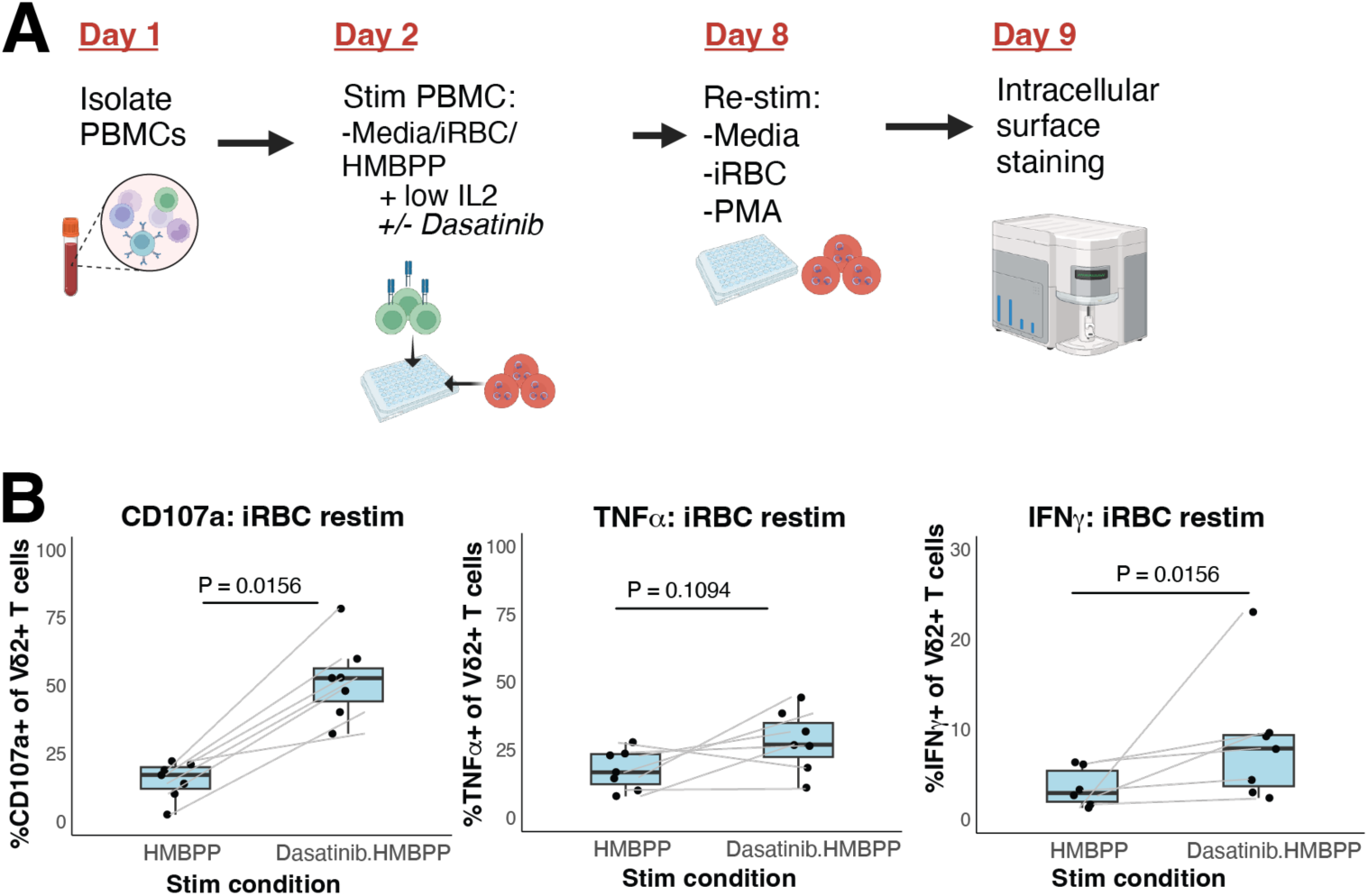
Diminished Vδ2+ T cell function following repeated malaria exposure can be regained after a period with limited malaria exposure. **A.** Schematic showing experimental design for *in vitro* stimulation experiments including dasatinib. Created in https://BioRender.com **B.** Vδ2+ T cell CD107a (left), TNFα (middle), and IFNγ (right) production in PBMC samples with HMBPP + Dasatinib compared to HMBPP alone. Values normalized to media re-stim condition.

## Discussion

Altogether, our data show that repeated *Plasmodium falciparum* exposure drives coordinated epigenetic and transcriptional reprogramming of circulating Vδ2+ γδ T cells that tracks with their diminished pro-inflammatory function. Across both a cross-sectional comparison of children living in high- (Tororo) *vs.* low-transmission (Jinja) settings and a longitudinal cohort spanning a marked decline in transmission following insecticide spraying, higher malaria exposure was associated with reduced Vδ2+ T cell TNFα and IFNγ regulation and with thousands of differentially accessible chromatin regions. Higher exposure consistently increased chromatin accessibility and/or expression at immunoregulatory and exhaustion-associated genes—including *LYN, PLCG2, HAVCR2* (Tim-3), *BATF, B3GAT1* (CD57), *CD8,* and multiple KIRs—whereas lower exposure was associated with increased accessibility and expression at pro-inflammatory and effector genes (*IFNG, TNF, GZMK, CD44, IL7R*) and at AP-1 (JUN/FOS) transcription factor motifs. cCRE-to-gene linkage analysis indicated that many of these loci, including *LYN* and *IL7R*, are coordinately regulated at the levels of chromatin accessibility and transcription. Single-cell profiling further identified a CD8+CD16+CD57+ Vδ2+ T cell subset enriched for this high-exposure signature (*LYN, PLCG2, HAVCR2*, KIRs) and reduced *IL7R*, consistent with a differentiated, chronically stimulated phenotype. Finally, an *in vitro* model of repeated stimulation recapitulated the hyporesponsive phenotype, demonstrating that reduced degranulation and cytokine production were predominantly TCR-dependent, accompanied by heightened p38 signaling, and partially reversible by dasatinib-induced TCR rest. Importantly, this hyporesponsive state was not fixed: pro-inflammatory function was regained as transmission and antigen exposure declined, both *in vivo* and *in vitro*.

Interestingly, Vδ2+ T cells following repeated malaria appear similar to γδ T cells expanded following tuberculosis (Mtb) infection (*28*). In the setting of chronic Mtb, γδ T cells were non-responsive to HMBPP and TCR signaling and upregulate CD8 and KIR receptors and have effector memory-like features. However, these Mtb-responsive cells skewed towards non-Vδ2+ subtypes, downregulated p38, and had increased cytotoxicity via CD16 (*28*). In our CITE-seq analysis, the same cluster that upregulated CD8 and CD57 also upregulated CD16, so it is possible that a CD16+ population was present in low abundance in our longitudinal analysis. Our results differ from a recent controlled human malaria infection (CHMI) study that showed accelerated activation and inflammation among Vδ2+ T cells responding to a second infection compared to a primary infection and a lack of any epigenetic regulation (*41*). This difference could be explained by the fact that children in Uganda experience higher parasite densities (leading to higher TCR stimulation) and many more than two malaria infections (leading to innate cells that are primed to respond unlike a malaria-naïve adult (*14, 21*)). Likely, children living in a malaria-endemic region still have responsive Vδ2+ T cells following a second infection and these cells only become hyporesponsive following chronic antigen exposure.

AP-1 signaling is complex and traditionally follows one of two pathways (*42*): 1) cytokines activate MAPK/p38 cascades, which then directly phosphorylates c-Jun (*43, 44*) or regulates AP-1 factors via phosphorylation of Sap-1a, 2) TCR mediated signaling upregulates c-Fos, then NFAT1/NFAT2, NFATc1 and IRF4, and leads to proliferation and cytokine production. P38 has 4 different isoforms (alpha, beta, gamma, delta) and the composition of these different isoforms can determine whether p38 signaling activates or inhibits AP-1 factors (*45*). Additionally, phosphorylation at c-Jun (whether performed directly or indirectly) can have very different downstream impacts on chromatin accessibility and transcription, depending on the specific site of phosphorylation and the presence of specific co-activators and/or co-repressors (*46*). Activation of Vδ2+ T cells *in vitro* by the phosphoantigens HMBPP or IPP lead to phosphorylation of p38 MAPK (as well as ERK1/2 and JNK), which then activates AP-1 factors and leads to production of pro-inflammatory cytokines (*34, 35*). Interestingly, p38 MAPK phosphorylation appears to be delayed in IPP stimulation compared with anti-CD3 stimulation (*47*). Consistent with our observations that p38 signaling increases and that accessibility at AP-1 factors decrease in Vδ2+ T cells in settings of high malaria, regulatory T cells and anergic CD4+ T cells expressed increased p38 activity (*48, 49*), and exhausted CD8+ T cells and anergic CD4+ T cells both express low levels of AP-1 factors (*50, 51*). There are at least 3 sites in c-Jun that when phosphorylated reduce DNA-binding affinity and therefore suppress transcription (*46*).

We identified *LYN* and *PLCG2* as both more accessible and more expressed at timepoints with higher malaria exposure. LYN is a tyrosine kinase that has well-characterized roles in B cell receptor (BCR)-induced signal transduction (*52*) and myeloid cells and clinical importance in autoimmune disease and many solid tumor cancers. T cells can be impacted by altered LYN expression in other cells; for example, in LYN-deficient mice, T cells produce more IFNγ due to higher BAFF production by myeloid cells (*53*), and regulatory T cell (Treg) populations are increased (*54*). LYN overexpression has been identified in two T cell subsets: SIRPα-expressing activated CD8+ T cells (*55*) and central memory T cells (*56*), the latter of which was hypothesized to negatively regulate TCR signaling. Dasatinib suppresses LYN activity in tumors and mast cells (*57, 58*). Interestingly, LYN expression in tumor cells was associated with poor survival overall but increased positive response to dasatinib in lung cancer patients, particularly non-smoker female patients with adenocarcinomas (*59*).

PLCγ2, which is encoded by *PLCG2*, plays an important role in B cell signal transduction and maturation, particularly BCR-induced JNK and p38 responses (*37*). PLCγ2 is also vital for NK cell cellular cytotoxicity (*60, 61*), indicating that it could play a similar role in other cell types. In T cells, it has been thought that PLCγ1 plays a much bigger role in TCR signaling and calcium mobilization. Recent data suggests, however, that PLCγ2 is required in primary T cells (*36*) and participates in TCR signal transduction and T-cell selection (*62*). While not much is known about PLCγ2 in γδ T cells, γδ T cells in tumor microenvironment of soft tissue sarcoma patients positively correlated with PLCγ2 expression and PLCγ2 expression further correlated with patient survival (*63*). While not the primary kinase responsible for PLCγ2 phosphorylation and activation in T cells, LYN can still contribute to PLCγ2 phosphorylation (*36*), so it seems likely that pathways activated in γδ T cells following malaria exposure activate both.

CD44 and IL7R pathways are less well characterized in γδ T cells but could feasibly be mechanisms of restored cytotoxic function following reduction in malaria exposure. CD44 can stimulate T cell signaling cascades via the TCR/CD3 complex, which leads to increased proliferation, c-Jun activation, and expression of activation markers (*64*). In mice, CD44+ γδ T cells produce IL-17A and protect against *Listeria monocytogenes (65)*, though in other infections including the parasite *Toxoplasma gondii*, CD44-induced cytokine production may contribute to immunopathology (*66*). Following viral infections, CD4+ and CD8+ T cells initially downregulate *IL-7R* (*33*), which is consistent with our observation that γδ T cells collected at high-exposure timepoints expressed lower *IL7R* compared to γδ T cells collected at low-exposure timepoints. Inαβ T cells, IL7R expression increases as part of formation of the memory response (*33*) and addition of recombinant IL-7 reduces T cell exhaustion and increases T cell proliferation (*67*); it is unknown whether similar patterns exist for γδ T cells.

Integrating these findings, we propose a model **(Fig 7)** in which repeated malaria drives Vδ2+ γδ T cells into a reversible, hyporesponsive, exhausted-like state through a LYN–PLCγ2–p38–AP-1 signaling axis that reshapes T cell receptor (TCR) responsiveness. We hypothesize that ongoing antigen exposure increases *LYN* accessibility and expression; LYN in turn phosphorylates PLCγ2, which activates JNK and p38 downstream of the TCR. Sustained p38 signaling phosphorylates c-Jun in a manner that reduces accessibility at AP-1 motifs and shifts the transcriptional program toward immunoregulatory genes, including HAVCR2 (Tim-3) and inhibitory KIRs. Because the cascade is initiated at a tyrosine kinase, it can be interrupted pharmacologically, consistent with the partial restoration of function we observed with dasatinib. Conversely, as antigen exposure wanes—whether naturally or following effective vector-control programs—increased CD44 expression and reduced p38 signaling lead to regained AP-1 accessibility and TCR signaling, reactivating pro-inflammatory and effector gene expression and increasing *IL7R* transcription. Increased IL7R expression may further promote IL-17 production via STAT3 (29). Together, this model reframes Vδ2+ T cell hyporesponsiveness not as fixed dysfunction but as a dynamic state tuned by the level of ongoing malaria exposure.

**Figure 7:**
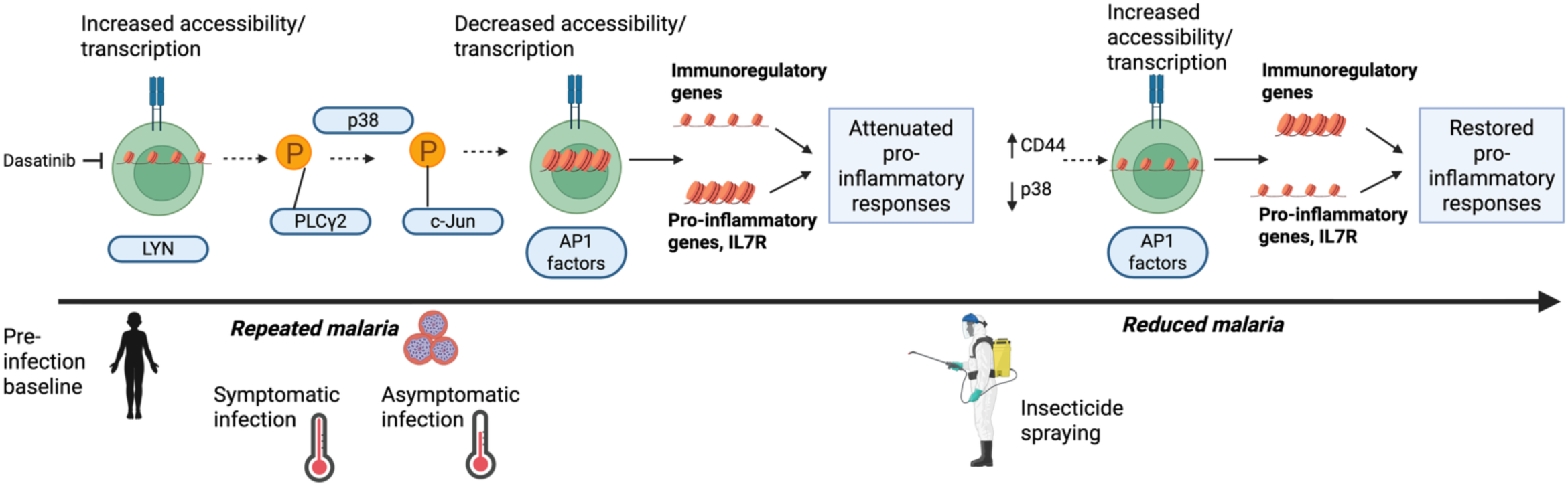
Model for possible mechanisms mediating altered Vδ2+ T cell function following repeated malaria. Created in https://BioRender.com

Our study has notable strengths and limitations. Its principal strength is the longitudinal sampling of the same children across a marked, programmatically driven decline in malaria transmission, which allowed us to track within-individual changes in Vδ2+ T cell function, chromatin accessibility, and transcription while minimizing inter-individual confounding. The integration of paired ATAC-seq, RNA-seq, and single-cell CITE-seq further enabled us to connect epigenetic and transcriptional states to defined cell subsets, and our *in vitro* stimulation model recapitulated key features of the *in vivo* phenotype while permitting mechanistic and pharmacologic testing. Several limitations should be considered. Our conclusions are largely associative: although our multi-omic data converge on a candidate LYN–PLCγ2–p38–AP-1 axis, we did not directly perturb these genes, and definitive causal attribution will require targeted genetic (e.g., CRISPR-Cas9) or pharmacologic approaches. Sample sizes were modest, and because annual entomological inoculation rate and calendar year were correlated, we could not fully disentangle cumulative exposure from time. It is also possible that insecticides directly modified Vδ2+ T cell transcription or function (*68*). We profiled circulating Vδ2+ T cells and therefore cannot speak to tissue-resident or Vδ1+ populations, and the timing of the observed epigenetic changes remain undefined. Finally, our *in vitro* model only partially reproduces chronic *in vivo* exposure, functional recovery with rest was incomplete and variable across donors, and the effects of dasatinib on iRBC-stimulated cells may be confounded by direct activity of the drug on the parasite.

Together, our findings suggest that repeated malaria reversibly reprograms the epigenome, transcriptome, and function of circulating Vδ2+ γδ T cells, and that reduced transmission can restore their pro-inflammatory capacity. These results position Vδ2+ T cells as modulators of clinical immunity to malaria, underscore the importance of monitoring the immunological consequences of malaria-control programs, and nominate the LYN–PLCγ2–p38–AP-1 axis as a candidate target for therapeutically restoring γδ T cell antimalarial function.

## Methods

### Ugandan cohort & sample collection

Samples were obtained from participants enrolled in PRISM1 or PRISM2 studies, part of the East Africa International Centers of Excellence in Malaria Research (U19 AI089674)(*24*). Since 2011, the ICEMR has followed children living in different malaria transmission settings in Uganda, including both high and low malaria transmission settings (*24*). In these settings, malaria transmission is year-round, with two seasonal peaks and varied transmission intensities. In Tororo District, before December 2014, malaria control was limited to the distribution of long-lasting insecticide-treated bednets (LLINs) and case management. IRS with the carbamate bendiocarb was first initiated in Tororo in December 2014, with additional rounds administered in June 2015 and November 2015. In June 2016, IRS was administered with the organophosphate pirimiphos-methyl (Actellic), with repeated rounds in June 2017 and June 2018. IRS was not initiated in Jinja District.

In both sites, all children from 100 randomly selected households were enrolled in the cohort study if they met the following eligibility criteria: (i) documented age between 6 months and less than 10 years, (ii) full-time resident of the household, (iii) no intention to move out of the subcounty for the next 2 years, (iv) agreement to come to a dedicated study clinic located within the subcounty for any febrile illness, (v) agreement to avoid antimalarial medications administered outside the study, and (vi) provision of written informed consent from parent or guardian (*29*). At enrollment, study participants and their parents/guardians were given an LLIN and underwent a standardized evaluation, including evaluation for any chronic medical conditions. No children were found to be HIV-infected.

Cohort participants are followed for all medical care at a dedicated study clinic open daily, undergo monthly routine visits for evaluation of parasitemia, and have banking of peripheral blood mononuclear cell (PBMC) at scheduled visits. Symptomatic malaria is defined as a documented fever (≥38.0°C) or history of fever in the previous 24 hours combined with a thick smear positive for malaria parasites. Participants with asymptomatic parasitemia were not treated with antimalarial drugs in accordance with local guidelines. Household aEIR in the year before sampling was estimated from monthly Centers for Disease Control and Prevention (CDC) light trap mosquito collections. The aEIR was the product of the annual human biting rate (total number of female *Anopheles* mosquitoes captured/number of house nights of collection × 365 days/year) and the sporozoite rate (number of mosquitoes testing positive for *Pf* sporozoites/number of mosquitoes tested). Clinical characteristics of the samples used can be found in **Table S1** (regional study) and **Table S3** (longitudinal study). Written informed consent was obtained from the parent or guardian of all study participants. The study protocols were approved by the Uganda National Council of Science and Technology (HS 1019), the Makerere University School of Medicine Research and Ethics Committee (2011-167), the University of California, San Francisco Committee on Human Research (11-05995), and the Institutional Review Board at Stanford University (41197).

### Production of frozen Plasmodium falciparum iRBCs

3D7 *Plasmodium falciparum* parasites were cultured in standard conditions and purified at schizont stages by MACS purification (Miltenyi Biotec, Auburn, CA). Parasites were frozen in 25ul human serum or fetal bovine serum (FBS) and 3x the volume of the iRBC pellet of glycerolyte to the pellet. To thaw iRBCs for stimulation experiments, 20ul of 3.5% NaCl was added to each tube and rested for 5 minutes. 1mL of the same solution was added, and the sample was centrifuged and the supernatant removed. After scraping the tube along the biosafety cabinet grate to dislodge parasites, the parasites were washed in 1.8% NaCl and then PBS. Parasites were counted using a hemocytometer.

### Bulk ATAC-seq

To probe the epigenetic landscape, we used protocols and analysis software for Fast-ATAC (*69*)—a fast and sensitive version of ATAC-seq (*23*) requiring only 5000 cells and optimized for blood cells. This method uses hyperactive Tn5 transposase, which inserts sequencing adapters into chromatin-accessible regions. In brief, cells were sorted into FACS buffer and pelleted by centrifugation at 500g RCF at 4°C in a pre-cooled fixed-angle centrifuge. The supernatant was removed carefully so as to not disturb the un-visible cell pellet. Fifty microliters of transposase mixture (25 ul of 2x TD buffer, 2.5 ul TDE1, 0.25 ul 2% digitonin, 22.5 ul nuclease-free water) was added to the cells. After mixing, samples were incubated at 37°C for 30 min in an Eppendorf ThermoMixer with agitation at 300 rpm. Following this transposition reaction, the transposed DNA was purified using a QIAGEN MinElute Reaction Cleanup kit (28204) and purified DNA was eluted in 10 ul elution buffer (10mM Tris-HCl, pH 8). Transposed fragments were amplified and purified and libraries quantified using qPCR before sequencing. Fast-ATAC libraries were sequenced using paired-end, dual-index library sequencing on an Illumina NextSeq instrument (75bp kit, 36 base pair reads, ∼20-30 million reads/sample).

### ATAC-seq analysis

The sequenced data were aligned to Human Genome Build hg38 and processed with an in-house SnakeATAC pipeline, using the Gencode v27 gene model (*70*). Downstream analyses of ATAC-seq data were conducted using the ChrAccR R package (*71*). Consensus peaks were called using MACS2, which were aggregated over regions (Tororo and Jinja) and the year of sampling (2013-16) for the Pilot and Longitudinal datasets, respectively. The agreement percentage was chosen such that at least 100,000 peaks were called **(Fig S1C** and **Fig S2B**). Samples below a fragment number of 100,000 were removed. Further samples were removed based on TSS enrichment < 3 and fragment number < 1000000 **(Fig S1B, Fig S2A**). Principal Component Analysis (PCA) was performed on quantile-normalized, log2-transformed consensus peaks.

Due to the small sample size and high intra-group variability of the regional dataset, ChromVAR differential analysis on JASPAR 2018 motifs was computed region-wise using limma::eBayes on scaled deviation scores (*72, 73*). Deviation scores for significant motifs (moderated t-tests using Limma;|log2FC| > 0.6, adjusted padj < 0.05) were plotted as a heatmap, generated with pheatmap. Donor-adjusted longitudinal data with Limma’s removeBatchEffect (70) was used for ChromVAR differential analysis with non-redundant transcription factor families (25). Given the larger cohort size of the longitudinal dataset, Wilcoxon rank-sum test was applied to compute p-values between groups of high and low Annual Entomological Inoculation Rate (aEIR, aEIR >= 100 vs. aEIR < 100), followed by multiple testing correction (Benjamini-Hochberg method). Differential motifs (Padj < 0.05, |delta| > 1) were plotted as a heatmap using pheatmap. Responder samples were identified from non-responders from within-group k-means clustering of differential motif families.

All differential analysis was performed using DESeq2 on the consensus peaks. The threshold for calling differential peaks was set at a Benjamini-Hochberg adjusted p-value of 0.05 and |log2FC| > 0.6. The Longitudinal dataset was restricted to “TruePairs” -- donors who had at least one sample in both the comparison groups of the measure under analysis. Donors were added as a covariate in the differential call for the longitudinal dataset.

The enrichment of motif families in differential peaks was calculated using Locus Overlap Analysis (LOLA)(*74*). Significantly enriched motif families (odds ratio > 0.3, q-value < 0.05) were plotted as bar plots with ggplot2. Footprinting plots of transcription factors were obtained from ChrAccR by merging samples by region for the pilot dataset and by aEIR for the longitudinal dataset.

### Bulk RNA-seq

RNA extraction and sequencing was performed at The Stanford Genomics. RNA was extracted using Qiagen miRNeasy kit catalog # 217084 following standard protocol. The Smarter Ultra Low Input RNA kit, from TakaraBio formerly Clontech (Catalog# 634848) was used to generate cDNA from of total 1 gram of RNA using the standard protocol. Amplified cDNA was purified using SPRI Ampure Beads from Beckman Coulter (Catalog # A63880). The quality and quantity were assessed using a High Sensitivity DNA chip on the Agilent 2100 Bioanalyzer. Illumina Nextra-XT kit (Catalog #FC-131-1024) was used to generate libraries from 150pg of amplified cDNA following standard protocol. Equal nanomole of uniquely indexed libraries was pooled and sequenced on NovaSeq6000 (2X150 pair-end).

### RNA-seq analysis

Sequencing reads from RNA-seq were processed using adapter trimming with SeqPurge (*75*) and the FastQC tool (version 0.11.7) (*76*). Pseudoalignments to the transcriptome (Gencode v27) were obtained using Kallisto (version 0.46.0). Differential analysis was performed on the gene level using the edgeR suite (*77*) after importing the data with tximport (*78*). No batch effects were observed. Only reads that mapped to protein-coding, TR_C/D/J/V, sense intronic, sense_overlapping, and lncRNA genes were taken for downstream analysis. Genes on sex chromosomes, as well as mitochondrial (MT), and ribosomal (RB) genes were thereby discarded. The dataset was normalized using trimmed mean of M-values TMM normalization, as recommended in the tximport manual. Principal Component Analysis (PCA) was computed using prcomp and plotted with ggplot2.

As described for the ATAC-seq analysis, the differential analysis for the RNA-seq dataset was carried out in the TruePair approach. The differentials were computed using edgeR’s Generalized Linear Model-Likelihood Ratio Test (glmLRT function), with donors as covariates. Differentially expressed genes (DEGs) (Padj <0.05, |log2FC| >= 0.6) were plotted as a heatmap using the log-normalized counts per million (CPM), generated with pheatmap. Gene Set Enrichment Analysis (GSEA) was computed using clusterProfiler and significantly enriched (p-value 0.05) immune-related pathways were plotted as dotplots.

Gene activity of consensus peaks was computed with ChrAccR’s getRBFGeneActivities function which utilizes counts weighted by Gaussian radial basis function (RBF). The resultant matrix was CPM-normalized, and limma::removeBatchEffect was applied to remove donor-specific effects. Wilcoxon rank-sum test was applied, and gene activities deemed significant (adjusted P-value < 0.05, |delta| > 0.6) were intersected with the RNA-seq differential expression results. A scatter plot using ggplot2 was generated of the two differential sets.

### Peak-to-gene analysis

The count matrices of the RNA-seq dataset and its corresponding ATAC-seq dataset were log2(CPM+offset) transformed and corrected for donor effects using limma’s removeBatchEffect (*79*). To reduce the burden of multiple testing, only the differentially accessible peaks and differentially expressed genes by aEIR group-based analysis were considered in the analysis. Candidate peaks located within ± 250k from the transcription start site (TSS) of a gene were selected for downstream analysis. Pearson correlation was computed between the accessibility of candidate peaks and the expression of their nearest genes. Linkages with a Pearson r > 0.3 were visualized as a heatmap using ComplexHeatmap. The genome tracks of the highly correlated linkages were plotted using pyGenomeTracks (*80, 81*).

### CITE-seq

6 PBMC samples from 3 Ugandan children (each at a 2013 and 2015 timepoint) were thawed and rested for 1 hour, and 1 x 10^6 cells were removed from each. γδ T cells were purified from the remaining cells using STEMCELL Technologies negative purification kit. The cells were then re-combined with the corresponding 1 x 10^6 PBMCs to create samples enriched for γδ T cells but still containing all cell types. Cells were then washed in cell staining buffer (BioLegend, San Diego, CA) and re-suspended in 50µL cell staining buffer. Five µL of Human TruStain FcX Blocking reagent (BioLegend) was added to each sample and incubated for 10 mins at 4°C. An antibody cocktail was prepared allowing for 1 µg of each TotalSeq B antibody (TotalSeq - B0066 anti-human CD7 Antibody, TotalSeq - B0582 anti-human TCR Vδ2 Antibody, TotalSeq - B088 anti-human CD279 (PD-1) Antibody, TotalSeq - B0159 anti human HLA-DR Antibody, TotalSeq - B0064 anti-human CD123 Antibody, TotalSeq -B0410 anti-human CD38 Antibody, TotalSeq - B0168 anti-human CD57 Recombinant Antibody, TotalSeq™-B0207 anti-human CD370 antibody, TotalSeq™-B0160 anti-human CD1c Antibody, TotalSeq™-B0053 anti-human CD11c antibody, Isotype control IgG1, Isotype control IgG2b) for each sample. 50µL mastermix was transferred to TotalSeq - B Human TBNK Cocktail tube (BioLegend) and mixed. Mix was then transferred to 10 Eppendorf tubes and spun at 14,000xg at 4°C for 10 minutes before adding to cells. The liquid was carefully pipetted out and added to cell suspensions. Cells were incubated for 30 minutes at 4°C. Cells were then washed x3 in cell staining buffer and re-suspended at a concentration of 1million/ml in cold PBS. Cells were filtered before being transferred to the Stanford Genomics Core for library preparation and sequencing. About 10,000 cells per sample were loaded onto the 10x Chromium controller. Gene expression libraries were prepared for each sample according to the manufacturer’s protocol (10x Genomics). Cell surface protein libraries were subjected to double the manufacturer’s recommended primer concentration and seven or eight amplification cycles during the sample index polymerase chain reaction (PCR) to reduce the likelihood of daisy chains forming. All libraries were sequenced using NovaSeq 6000 (Illumina) to achieve a minimum of 50,000 paired-end reads per cell for gene expression and 20,000 paired-end reads per cell for cell surface protein libraries.

### Data processing for CITE-seq Analysis

Raw CITE-seq data were processed using Cell Ranger (v6.0.0, 10x Genomics), followed by ambient RNA correction using SoupX (v1.6.2). Cells were excluded if they contained fewer than (10^2.9) unique detected genes (nFeature_RNA), fewer than (10^3.4) RNA total RNAs (nCount_RNA), more than 12% mitochondrial transcripts (percent.mt), or fewer than (10^2.45) total ADT counts (nCount_Protein). Thresholds were determined by the distributions of these quality-control metrics across samples **(Fig S7A)**. More than 90% of cells were retained after quality-control filtering in all samples except 3089-1, which retained 85% of cells. Application of these thresholds yielded a final dataset of 30,373 high-quality cells across six samples for downstream analyses. Doublets were subsequently identified and removed using DoubletFinder (v2.0.3). Cell type annotation was performed using SingleR (v2.4.1) with the Monaco Immune Data reference atlas (*82*). Out of all cells from various cell types, γδ2 T cells were identified using expression of the γδ2 ADT marker. The distribution of CLR-normalized γδ2 protein expression across all cells exhibited a bimodal pattern (**Fig S8B**). Inspection of RNA-based SingleR annotations across expression bins demonstrated that the high-expression population was enriched for Monaco-annotated γδ2 T cells, whereas intermediate-expression cells represented a mixture of γδ2 and non-γδ2 populations. Based on these observations, cells with CLR-normalized γδ2 expression greater than 1.2 were selected for downstream analyses. For the transcriptomic assay, gene variance was calculated on log1p-transformed counts to exclude low-variance features and ribosomal protein genes, retaining the top 85% most variable non-ribosomal features. Dataset features were restricted to protein-coding genes and long non-coding RNAs (lncRNAs) mapped via annotations from Ensembl BioMart (hsapiens_gene_ensembl, version 114). RNA data were normalized using Seurat’s LogNormalize method, whereas ADT data were normalized using centered log-ratio transformation. Following scaling, principal component analysis was performed separately for RNA and ADT modalities. Sample-level batch effects were corrected using Harmony (v1.2.3), and UMAP embeddings were generated from the Harmony-corrected spaces. Residual non-γδ cells remaining after ADT thresholding were identified using the ADT principal component space in conjunction with Monaco-based annotations and excluded prior to downstream analyses (**Fig S8C**), and individual cells falling outside the RNA principal component boundaries (**Fig S8D**) were removed. Cellular clustering was executed by constructing Shared Nearest Neighbor (SNN) graphs and applying the Louvain community detection algorithm across multiple resolutions ranging from 0.3 to 1.4 for both assays. For the ADT modality resolution of 0.3 was chosen while for the RNA modality a resolution of 0.5 was chosen (**Fig S8E**). Differential expression analyses between specific cell clusters and background populations were conducted utilizing the Wilcoxon rank-sum test, incorporating a threshold of detection in at least 10% of cells, an adjusted p-value threshold of less than 0.05, and a cut-off of 0.6 absolute log2 fold-change. To evaluate concordance with bulk transcriptomic signatures, single-cell log fold-change (logFC) estimates were directly compared against matched bulk RNA-seq differential expression results using scatter plots. Gene Ontology (GO) Biological Process enrichment analyses on concordant and direction-specific marker gene sets were performed using clusterProfiler and org.Hs.eg.db, utilizing Benjamini-Hochberg adjustment for multiple testing with significance cutoffs of *p* < 0.05 and *q* < 0.05.

### In vitro *stimulation assay*

Leukocyte reduction system (LRS) chambers were obtained from the Stanford Blood Bank. PBMCs were isolated using Ficoll and cells were left to rest overnight in R10 Media (RPMI supplemented with 10% FBS, Pen-Strep, L-glutamine and HEPES). For experiments using purified γδ T cells, PBMC were washed in MACS buffer (PBS containing 10% FBS and 0.5M EDTA) and γδ T cells were isolated using a negative purification kit from STEMCELL Technologies (Vancouver, Canada). Cells were counted on a Countess Cell Counter (Thermo Fisher Scientific, Waltham, MA) and set up for stimulation assays in 96-well plates with 250,000 – 500,000 cells/well in R10 media made with human serum and containing 30 IU IL-2 (to ensure high cell viability for the purified γδ T cell conditions). Infected red blood cells (iRBC) were thawed as described above and added at a ratio of 3:1 iRBC:cells. HMBPP (1-Hydroxy-2-methyl-2-buten-4-yl 4-diphosphate, Echelon Biosciences, Salt Lake City UT) was added at a final concentration of 10 nM. Cells were spun down and supplemented with fresh media after 3 days. After 6 days (or following 3-5 days rest following 6-day stim for experiments testing rest conditions), cells were counted and re-stimulation assays were set up with 100,000 cells/well and iRBCs added at a ratio of 1:1. After 6 hours, brefeldin/monensin (GolgiStop/GolgiPlug, BD Biosciences, Franklin Lakes, NJ) and APC-conjugated anti-CD107a (BioLegend 328619, clone H4A3, 0.8 µl/well) were added to all media and iRBC wells. The following morning, PMA (Sigma-Aldrich), Ionomycin (Sigma-Aldrich), brefeldin/monensin, and CD107a were added to all positive control wells and left in the incubator for 4-5 hours before staining. For experiments using dasatinib, dasatinib (Sigma-Aldrich, St. Louis, MO) was added at (1 µM) (*39*) during the 6-day stimulation period.

### Flow cytometry

Cells were washed in FACS Buffer (PBS containing 0.5% bovine serum albumin and 2% 500 mM EDTA) and then incubated in surface antibodies (see table below) in PBS for 30 minutes at room temperature. Cells were washed twice and then incubated in 40 µL FixA solution (Thermo Fisher Scientific, Waltham, MA) for 15 minutes. Following 1 wash, cells were incubated in intracellular cytokine antibodies (TNFα BV421 BioLegend 502932, clone MAb11; IFNγ BV785 BioLegend 502542, clone 4S.B3) suspended in PermB (Thermo Fisher Scientific) for 30 minutes. Finally, cells were washed x3, re-suspended in PBS, and run on a Attune Nxt Cytometer (Thermo Fisher Scientific). Compensation was performed using UltraComp beads (ThermoFisher Scientific) with single stains added. Paired non-parametric (Wilcoxon ranked sum) tests were used to compare expression between stimulation conditions. Data was visualized using ggplot2 or Graphpad Prism.

*Surface antibodies – italicized ones used for all experiments*

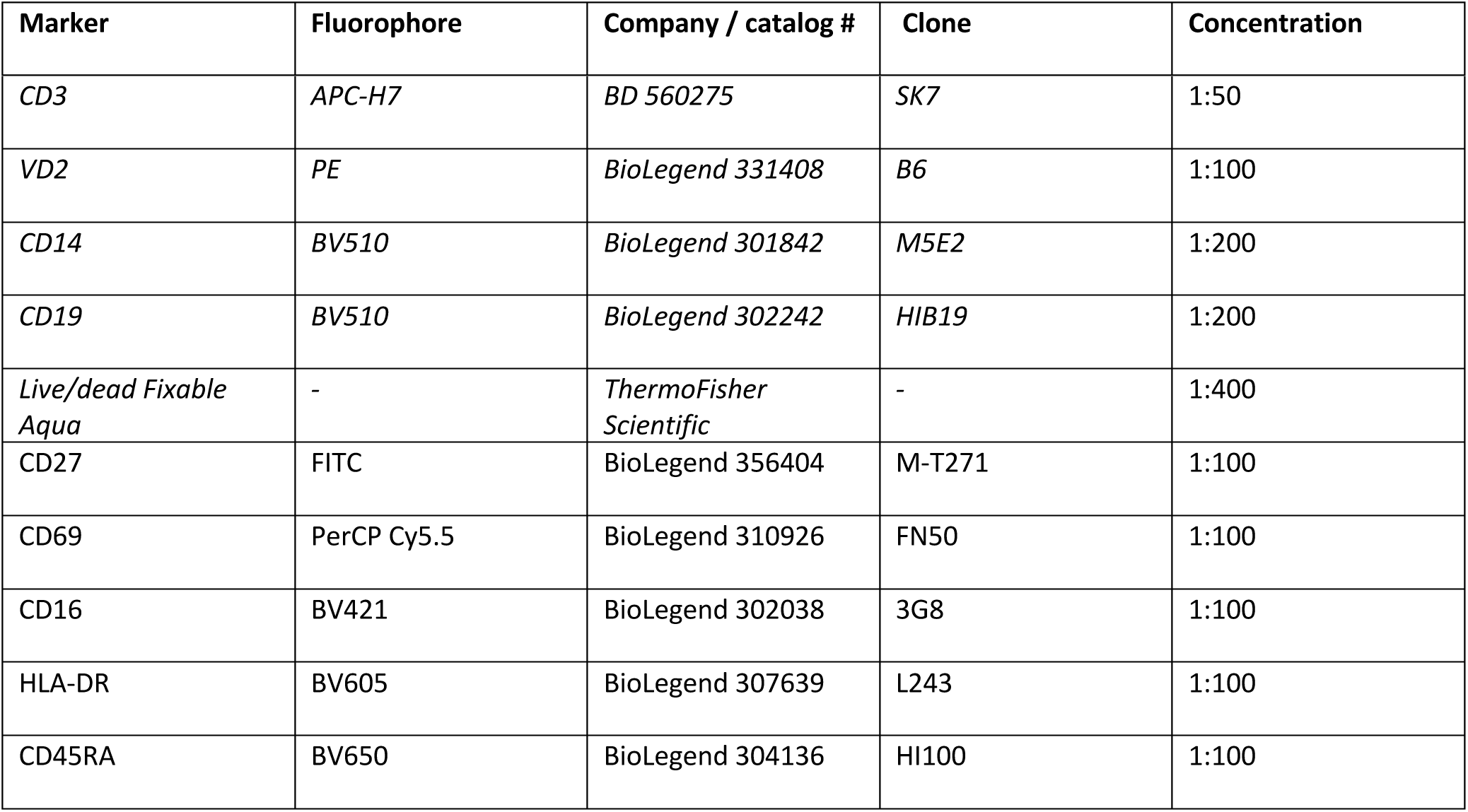

### Phosphoflow

Cells were washed in FACS Buffer and then incubated in surface antibodies (see list below + italicized from previous section) in PBS for 30 minutes at room temperature. Cells were stimulated for 1-32 minutes with iRBC, HMBPP or PMA/Ionomycin or Pervanadate controls. Following stimulations, cells were immediately mixed with warm Phosflow Cytofix buffer (BD Biosciences) and incubated at 37°C for 10 minutes. Following centrifugation and removal of supernatant, cells were permeabilized in cold Phosflow Perm Buffer III (BD Biosciences) and incubated for 30 minutes on ice. Cells were washed x3 and incubated with phospho antibodies for 1 hour. Cells were washed x3, re-suspended in PBS, and run on a Attune Nxt Cytometer (Thermo Fisher Scientific).

*Phosphoflow antibodies:*

- Phospho-PLC-gamma-1 (Tyr783) (D6M9S) Rabbit mAb (Alexa Fluor(R) 488 Conjugate) - Cell Signaling Technology 25678S, 1:50
- Phospho-p38 MAPK (Thr180, Tyr182) Monoclonal Antibody (4NIT4KK), PE, eBioscience™ (Thermo Scientific 12-9078-41), 1:10
- Phospho-ZAP70/Syk (Tyr319, Tyr352) Monoclonal Antibody (n3kobu5), PerCP-eFluor 710, eBioscience™ (Thermo Scientific 46-9006-41), 1:10
- Erk1/2 (pT202/pY204) (p44/42 MAPK; Extracellular signal Regulated Kinase 1/2 (pT202/Y204)) Mouse, Alexa Fluor 647, Clone: 20A, BD 612593, 1:2.5

### ADCC

PBMCs or purified γδ T cells were prepared at 4 million cells/mL in 100 µL R10 media. Infected red blood cell (iRBC) opsonization was performed by adding 10% Ugandan or American serum or anti-RBC positive control (Abcam ab34858) to 5 million iRBCs in 100 µL incomplete RPMI media. Tubes were incubated at 37°C for 1 hour, with mixing every 15 minutes. Cells were washed 1.5x, re-suspended in R10 media, and plated in a 96-well plate. PBMCs or purified γδ T cells at an effector:target ratio of 1:2 (i.e. 500,00 PBMC: 1x10^6 iRBC). Brefeldin/monensin (GolgiStop/GolgiPlug, BD Biosciences) and APC anti-CD107a (0.8 µl/well) were added to all wells, and the plate was spun for 3 minutes at 100g. Cells were then incubated at 37°C for 5 hours and staining for flow cytometry was performed as mentioned above. Antibodies used included those italicized in “surface antibody” table, as well as CD16 BV711 (BioLegend 302044, clone 3G8).

## Data and code availability

For the ATAC-seq/RNA-seq dataset, sample annotation, peak-level accessibility quantifications, ChromVAR quantifications, and gene-level expression are available on GEO. Scripts used for all the analyses are publicly available and can be found on GitHub (https://github.com/EpigenomeInformatics/vd2_malaria) under MIT license.

## Funding

This study received financial support from the National Institutes of Health (U19 AI089674 (GD, MK, MF), U01AI150741 (BG, IRB, PJ); the Gates Foundation INV-052649 (PJ, JN); and the Stanford Maternal and Child Health Research Institute, Woods Family Faculty Scholar in Pediatric Translational Medicine (PJ). KDP was supported by AP Giannini Fellowship and Stanford Maternal and Child Health Institute postdoctoral fellowship. An ASTMH Travel Fellowship funded some experiments in Uganda. RNA sequencing data was generated with instrumentation purchased with NIH funds (award numbers: S10OD025212 and 1S10OD021763).

## Acknowledgements

Cell sorting was done on instruments in the Stanford Shared FACS Facility (RRID: SCR_017788). RNA-seq and CITE-seq experiments were performed in the Stanford Genomics Core facility. CYTOF experiments were done in Human Immune Monitoring Center with support from Michael Leipold, Gerlinde Obermoser, and Holden Maecker.

## Author contributions

KDP and PJ conceptualized the project. SK, FM and PJ supervised the project. MIJM developed computational pipelines for analysis of bulk ATAC-seq and RNA-seq data, performed associated statistical analysis with support from FM, SK, and J S-H, and made figures associated with this analysis. MIJM curated the datasets and managed GEO submissions. AAM performed all analysis of CITE-seq data with support from FM and SK, and made figures associated with this analysis. SK and WG provided additional guidance on ATAC-seq, RNA-seq, and CITE-seq. KDP, KC, and MB performed *in vitro* stimulation experiments; KDP made associated figures. DC and AK processed ATAC-seq samples. VS processed RNA-seq samples. MT and JN processed CITE-seq and CYTOF samples. JR, FN, IS, MK, BG, GD and MF supervised collection of samples in Uganda. J S-H, BG, GD and MF provided funding. KDP made the final figures and wrote the manuscript. All authors read the manuscript and provided feedback.

## Notes

### Competing Interest Statement

The authors have declared no competing interest.

