## Supplemental Figures for "Repeated malaria exposure reprograms Vδ2+ γδ T cell chromatin accessibility and immune function toward a reversible hyporesponsive state"

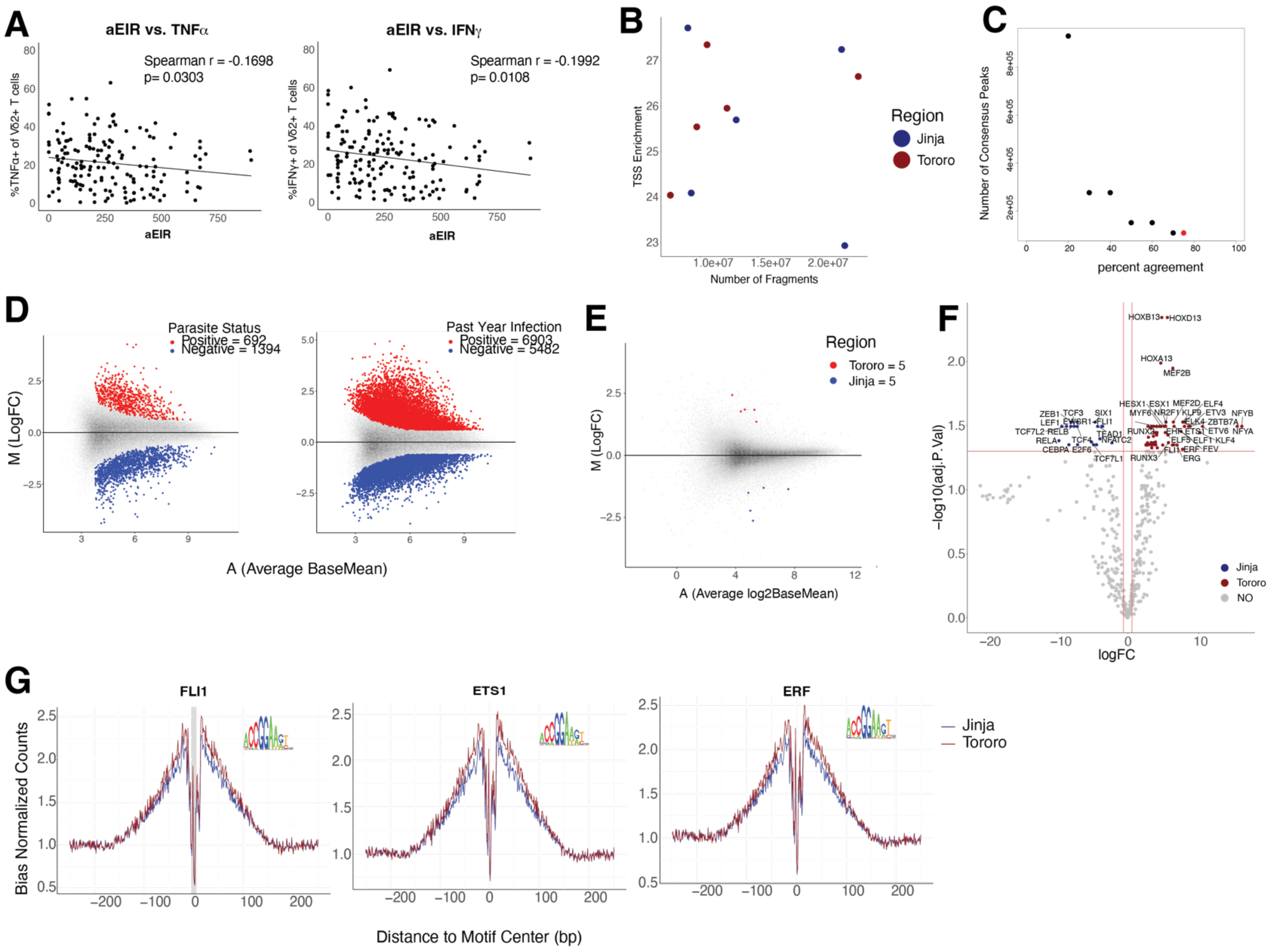


**Figure S1: Additional analysis of differential cytokine responses, chromatin accessibility and transcription factor binding among samples from high vs. low transmission regions**

**A.** PBMC from 155 children in Tororo (highly endemic region) and 8 children living in Jinja (low transmission) were stimulated with iRBC for 24 hours. Production of TNFα and IFNγ by Vδ2⁺ T cells in response to iRBC were significantly correlated with aEIR.

**B.** Scatter plot of samples' (colored by region) fragment number against TSS enrichment.

**C.** Dot plot illustrating the percent agreement by region for defining consensus peaks using MACS2. The red point indicates the selected fragment agreement threshold.

**D.** MA plots highlighting differentially accessible regions for Parasite Status (positive vs negative; N=3 positive and 7 negative) and Past-year Infection Status (positive vs. negative; N=4 positive and 6 negative) (|logFC| > 0.6, FDR < 0.05).

**E.** MA plot showing differentially accessible regions for CD4 T cells

**F.** Volcano plot of ChromVAR DARs by region

**G**. Transcription Factor Footprint plot of FLI1, ETS1, and ERF, showing more accessibility in Tororo (high malaria exposure) around TF binding sites.

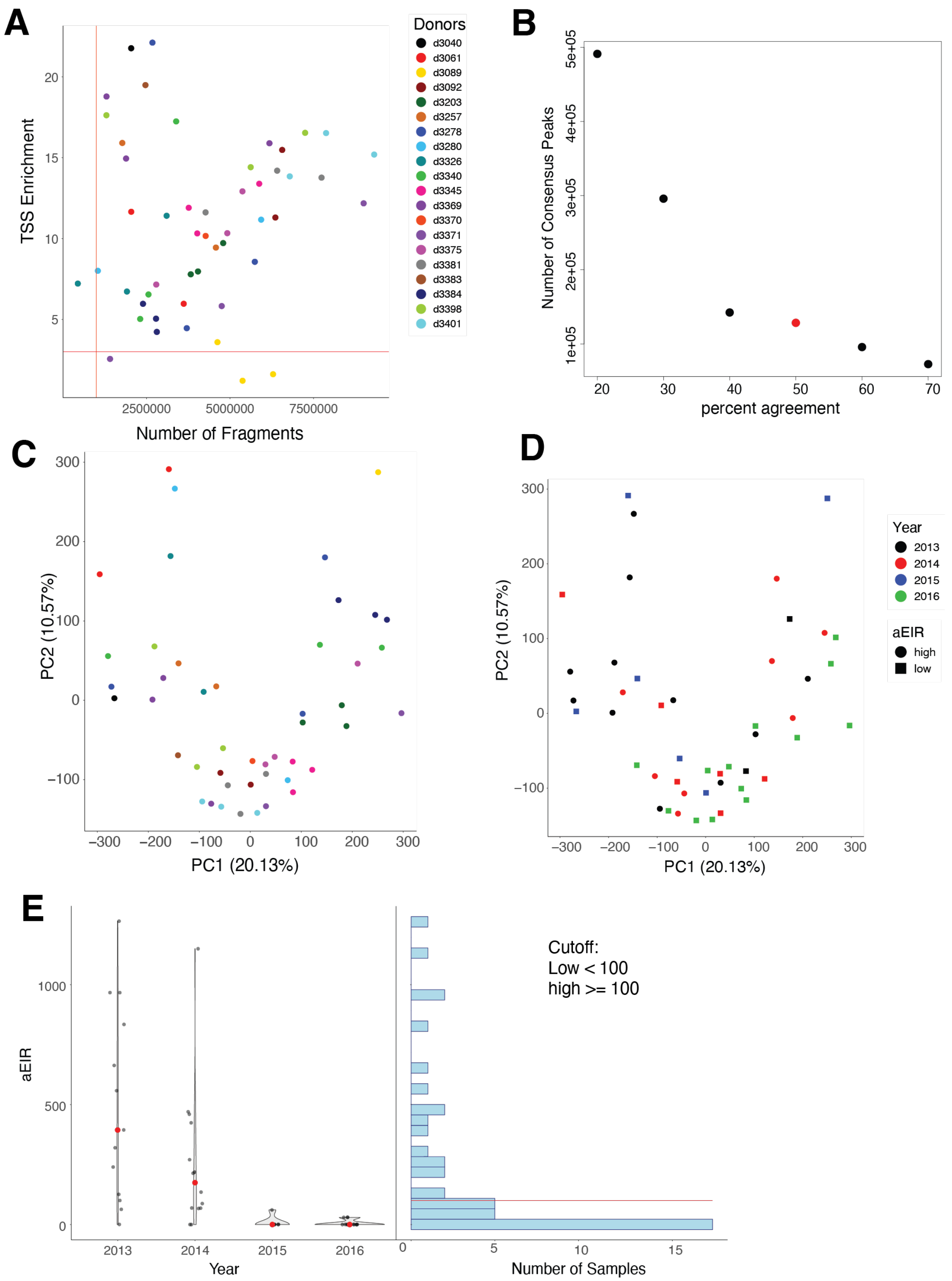


**Figure S2: Quality control and distribution of longitudinal samples for ATAC-Seq**

**A.** Scatter plot of samples' fragment number against TSS enrichment. The red lines denote the cutoff set for QC.

**B.** Dot plot denoting the Percentage of fragments showing agreement by year measure (2013 – 2016) to call a consensus peak. Red dot denotes the percentage agreement of the data used for analysis.

**C-D.** PCA of the samples by donors (C) and year/aEIR (D).

**E.** Violin plot illustrating the relationship between year and aEIR (left) and histogram of aEIR counts (right). The red line represents the threshold used to group the samples into low and high categories.

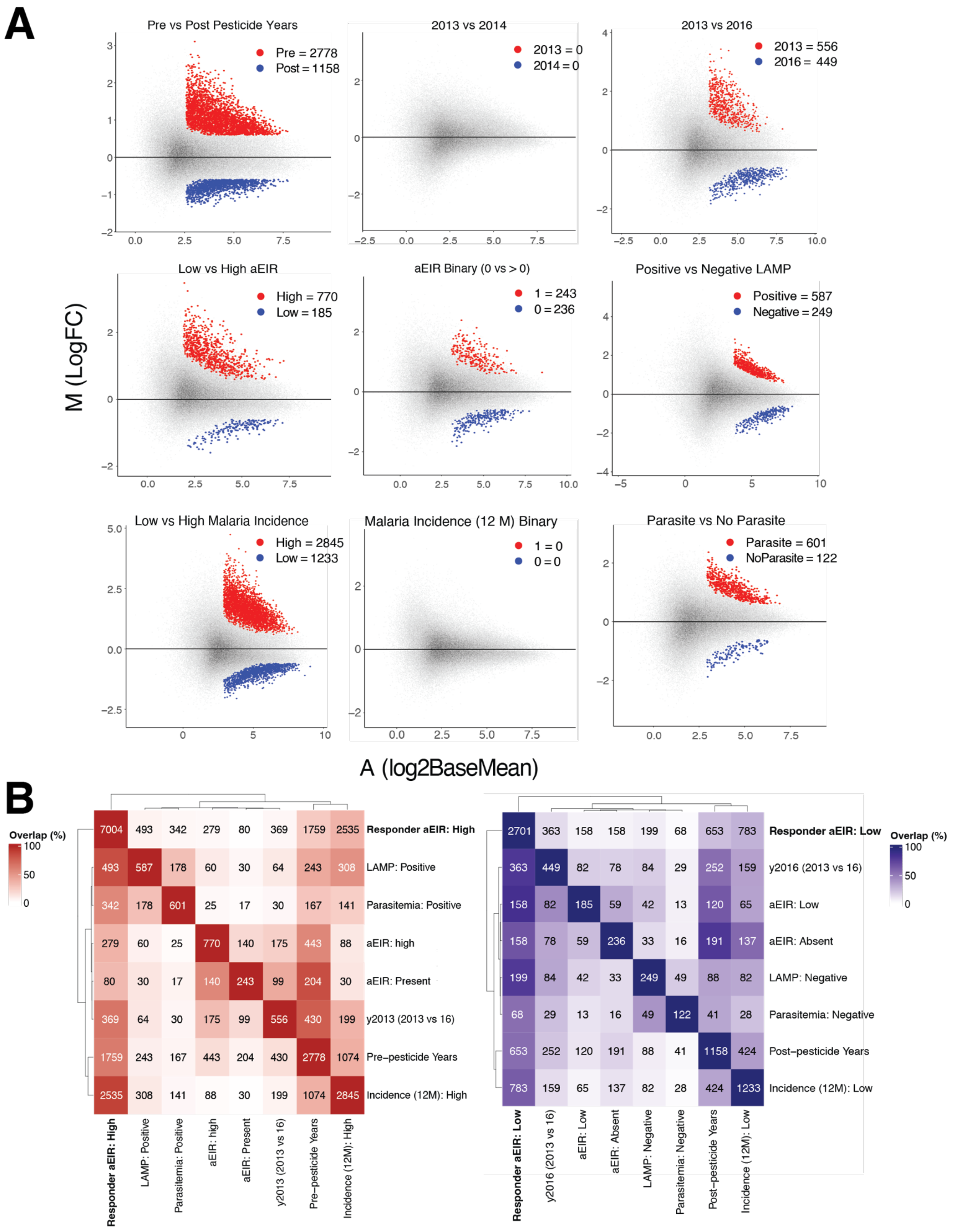


**Figure S3: Relationship between differentially accessible regions for different clinical or exposure features**

**A.** Matrix of DARs across different measures for the consensus peaks across all samples (responders as well as non-responders). Pre pesticide=2013 or 2014 (N=25). Post pesticide=2015 or 2016 (N=15). For 2013 vs. 2014, N= 11 for each. For 2013 vs. 2016, N=10 for each. Low EIR <100 (N=14). High EIR >100 (N=19). For aEIR binary comparison, N=10 for aEIR=0 and 15 for aEIR>0. LAMP= parasite detection by Loop-Mediated Isothermal Amplification (includes submicroscopic; N=5 positive and 5 negative). Incidence=number of symptomatic infections within previous year. Low incidence= 0-1 infections (N=7). High incidence = 2-7 infections (N=9). For binary incidence comparison, N=14 for each group. Parasite vs. no parasite= any microscopic or submicroscopic parasites detected (N=11) vs. not detected (N=10).
**B.** Row-scaled confusion matrix denoting the percentage of peaks overlapping with the DARs in (A) and Fig 2D. Numbers within the tiles indicate the absolute number of overlapping peaks. Color scheme reflects the percentage of overlaps.

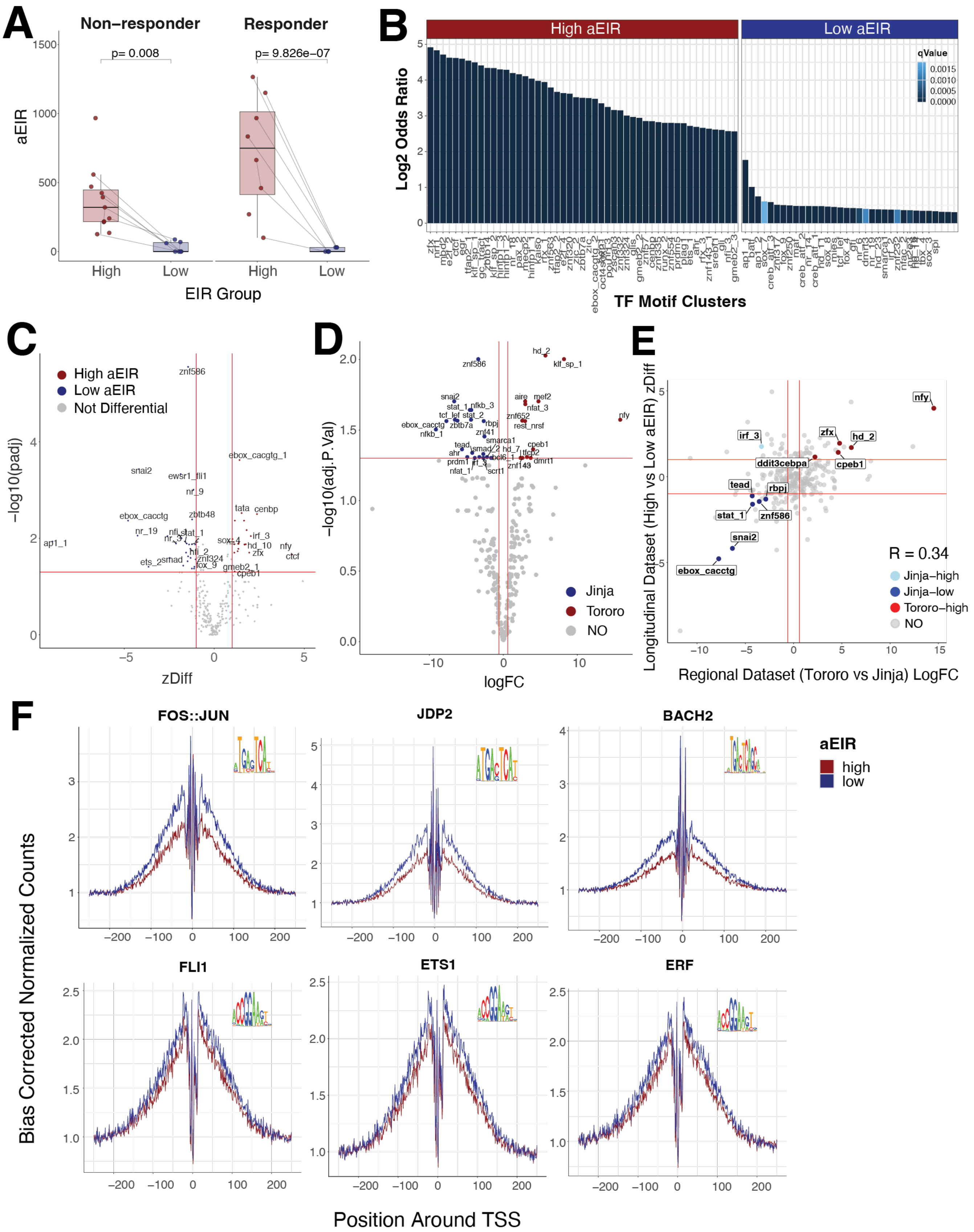


**Figure S4: Comparison of transcription factor binding motifs for longitudinal vs. cross-sectional studies**

**A.** Comparison of aEIR differences between High and Low groups for responder samples (clusters 1 and 3 in Fig 2C) and non-responders (cluster 2).
**B.** Bar plots showing motif families enriched in longitudinal LOLA analysis (qval < 0.05, |logOR| > 0.3)
**C.** Volcano plot of longitudinal data set's differential motif activity (ChromVAR) by aEIR
**D.** Volcano plot of cross-sectional data set's differential motif activity (ChromVAR) by region
**E.** Scatterplot of the agreement of differential TF motif activity (ChromVAR) from C and D
**F.** Transcription Factor (TF) footprint plots for longitudinal samples. All TF binding sites shown demonstrate more accessibility at low aEIR timepoints

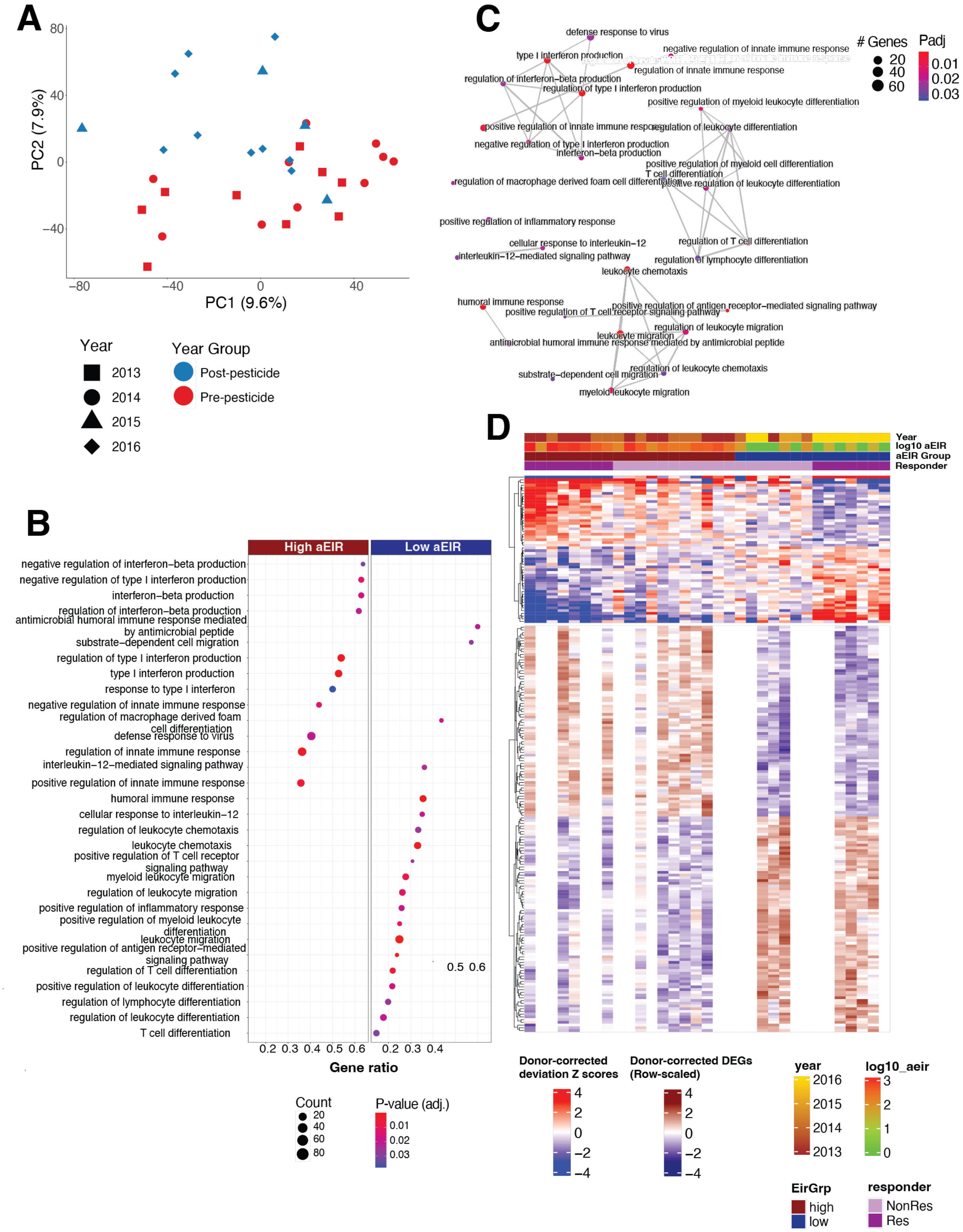
A
PC1 (9.6%)
PC2 (7.9%)
Year
2013
2014
2015
2016
Year Group
Post-pesticide
Pre-pesticide
B
High aEIR
Low aEIR
negative regulation of interferon−beta production
negative regulation of type I interferon production
interferon−beta production
regulation of interferon−beta production
antimicrobial humoral immune response mediated by antimicrobial peptide
substrate−dependent cell migration
regulation of type I interferon production
type I interferon production
response to type I interferon
negative regulation of innate immune response
regulation of macrophage derived foam cell differentiation
defense response to virus
regulation of innate immune response
interleukin−12−mediated signaling pathway
positive regulation of innate immune response
humoral immune response
cellular response to interleukin−12
regulation of leukocyte chemotaxis
leukocyte chemotaxis
positive regulation of T cell receptor signaling pathway
myeloid leukocyte migration
regulation of leukocyte migration
positive regulation of inflammatory response
positive regulation of myeloid leukocyte differentiation
leukocyte migration
positive regulation of antigen receptor−mediated signaling pathway
regulation of T cell differentiation
positive regulation of leukocyte differentiation
regulation of lymphocyte differentiation
regulation of leukocyte differentiation
T cell differentiation
Gene ratio
Count
P-value (adj.)
C
### Genes
Padj
defense response to virus
type I interferon production
negative regulation of innate immune response
regulation of innate immune response
regulation of interferon−beta production
regulation of type I interferon production
positive regulation of myeloid leukocyte differentiation
positive regulation of innate immune response
regulation of leukocyte differentiation
negative regulation of type I interferon production
interferon−beta production
positive regulation of myeloid cell differentiation
T cell differentiation
regulation of macrophage derived foam cell differentiation
positive regulation of leukocyte differentiation
positive regulation of inflammatory response
cellular response to interleukin−12
regulation of T cell differentiation
interleukin−12−mediated signaling pathway
regulation of lymphocyte differentiation
leukocyte chemotaxis
humoral immune response
positive regulation of antigen receptor−mediated signaling pathway
positive regulation of T cell receptor signaling pathway
regulation of leukocyte migration
leukocyte migration
antimicrobial humoral immune response mediated by antimicrobial peptide
regulation of leukocyte chemotaxis
substrate−dependent cell migration
myeloid leukocyte migration
D
Year
log10 aEIR
aEIR Group
Responder
Donor-corrected deviation Z scores
Donor-corrected DEGs (Row-scaled)
year
log10_aeir
EirGrp
high
low
responder
NonRes
Res

**Figure S5: Analysis of longitudinal RNA-Seq samples and association with motif activity**

**A.** Principal component analysis (PCA) plot of RNA cohort samples. PC2 variance is explained by the year (2013/2014 vs. 2015/2016).

**B.** Gene Set enrichment analysis (GSEA) of DEGs by aEIR (High vs Low).

**C.** Enriched Gene Ontology (GO) terms associated with the DEGs. Node size represents the number of enriched genes, and edge width corresponds to the degree of gene overlap between terms (calculated with Jaccard index).

**D.** Heatmaps of ChromVAR's motif activity (ATAC) and DEGs (RNA) showing concurrent increases in accessibility and expression between the high and low aEIR groups.

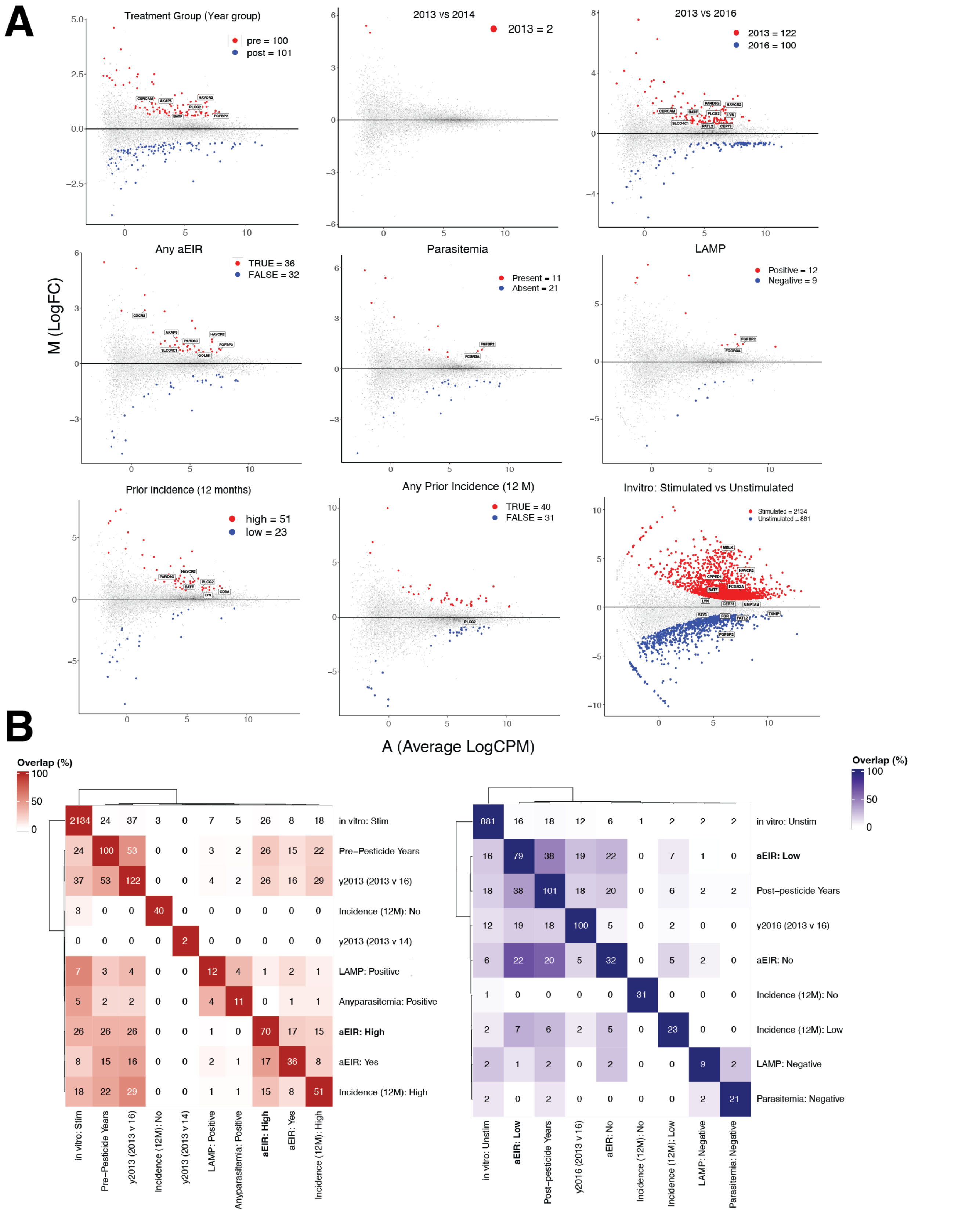


**Figure S6: Relationship between differentially expressed genes for different clinical or exposure features**

**A.** Matrix of differentially expressed genes for different measures for the same gene set. Pre pesticide=2013 or 2014 (N=17). Post pesticide=2015 or 2016 (N=11). For 2013 vs. 2014, N= 6 for each. For 2013 vs. 2016, N=6 for each. For aEIR>0, N= 8; for aEIR=0, N=6. Parasitemia = any microscopic or submicroscopic parasites detected

(N=7 positive and N=6 negative). LAMP= parasite detection by Loop-Mediated Isothermal Amplification (includes submicroscopic; N=2 positive and N=3 negative). Incidence=number of symptomatic infections within previous year. Low incidence= 0-1 infections (N=4). High incidence = 2-7 infections (N=6). For binary incidence comparison, N=3 for each group. *In vitro* stimulation= iRBC or HMBPP (N=2). Unstimulated N=1.

**B.** Row-scaled confusion matrix denoting the percentage of differential genes overlapping with the differentials in (A) and Fig 3A. Numbers in the tiles indicate the absolute number of peaks overlapping. Color scheme is based on percentage of overlaps.

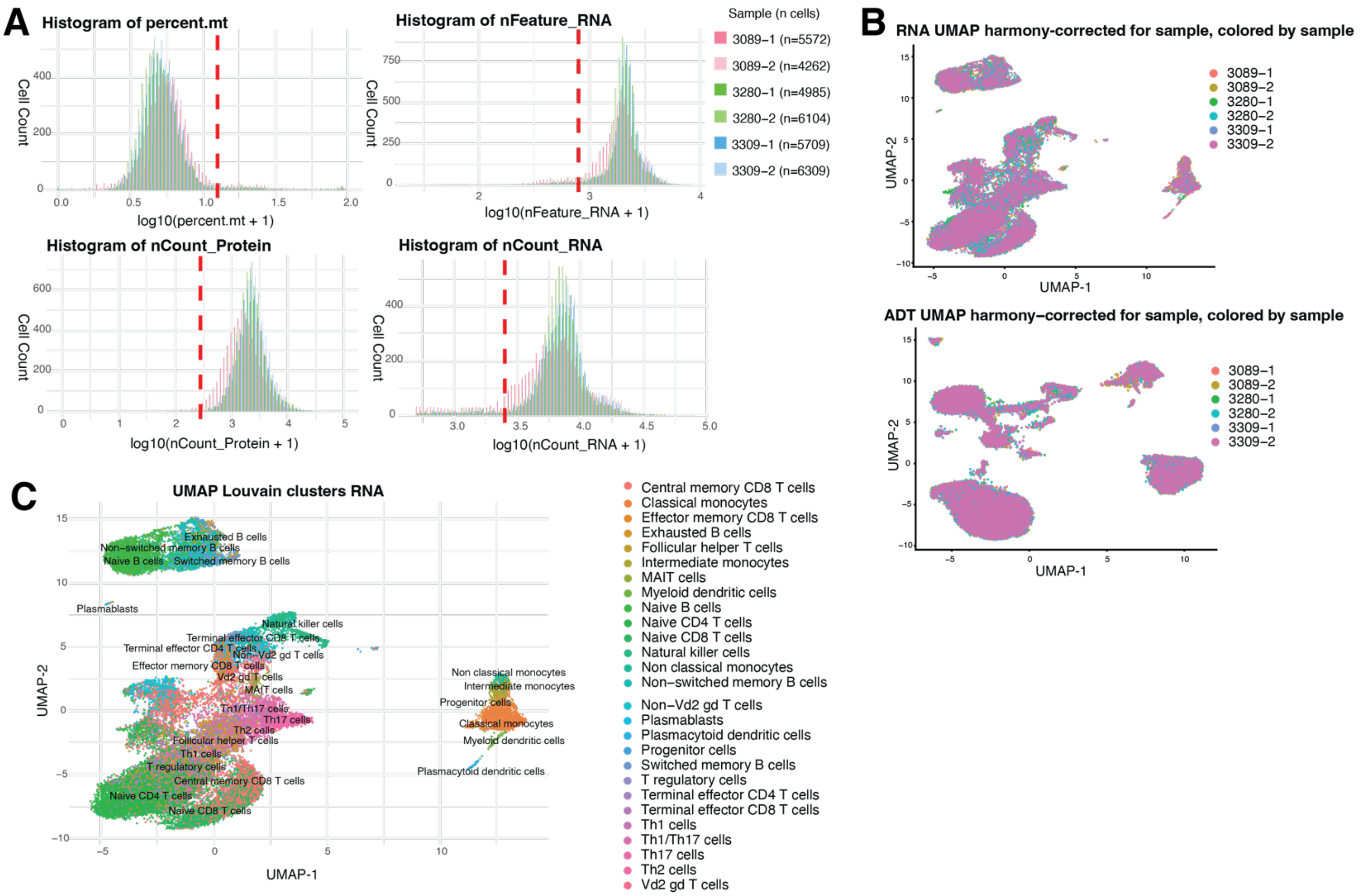


**Fig S7: CITE-Seq preprocessing and quality control**

**A.** A combined histogram showing RNA counts (nCount_RNA), RNA features (nFeature_RNA; number of unique genes detected per cell), protein counts (nCount_Protein), and mitochondrial transcript percentage (percent.mt), and supporting the application of global filtering thresholds. Metrics are log-transformed (log10 + 1) to account for skewed distributions. Samples are color-coded with cell counts indicated in the legend, and red dashed vertical lines mark thresholds used to filter out low-quality or dead cells during quality control (nFeature_RNA $\geq 10^{2.9}$, nCount_RNA $\geq 10^{3.4}$, nCount_Protein $\geq 10^{2.45}$, percent.mt $\leq 12\%$).

**B.** UMAP embeddings for RNA and ADT expression layers reveal differences in clustering patterns between samples.

**C.** UMAP projection (batch corrected for the sample batch variable using Harmony) colored with RNA-reference-based annotation using the Monaco immune atlas for cell-type assignment.

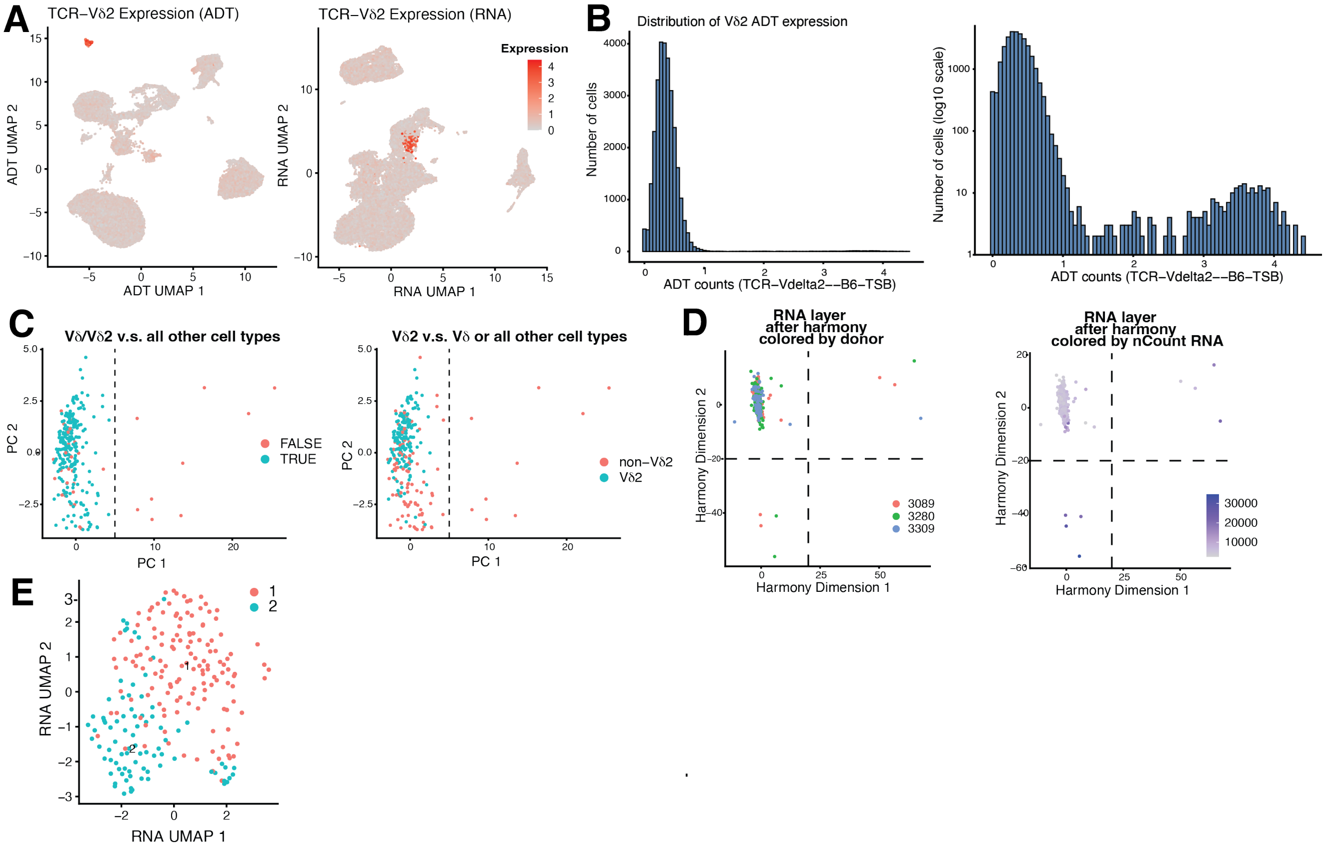


**Figure S8: Identification of Vδ2+ T cell populations from CITE-seq**

**A.** Vδ2+ T cells clustered by surface marker abundance (left), and in gene expression space (right) Clusters were Harmony corrected for the sample batch variable and cells are colored with the expression of the Vδ2 ADT.

**B.** Histogram showing the overall bimodal distribution of antibody-derived tag (ADT) expression counts for the Vδ2 TCR chain (TCR-Vdelta2--B6-TSB) across all sequenced cells, displayed on a $\log_{10}$ scale. The distinct right-hand peak corresponds to the targeted Vδ2+ population.

**C.** Principal component analysis (PCA) based on ADT expression. Left panel illustrates the separation threshold (vertical dashed line) used to refine the true Vδ/ Vδ2+ target population based on the Monaco immune atlas annotation (blue) from all other baseline cell types (red). Right panel demonstrates the isolation of the strict Vδ2+ subpopulation (cyan) against non-Vδ2+ cells (red). This ADT-based dimensionality reduction further separated Vδ2+ T cells from non-target T-cell populations and cells with ambiguous TCR signal and Monaco annotation consensus, yielding a high-confidence Vδ2+ T-cell subset for downstream analyses.

**D.** Harmony-based dimensionality reduction based on RNA expression. The scatter plot illustrates the separation thresholds (dashed lines) used to refine the true Vδ2+ target population from outlying cells across donors. Left panel shows coloring by sample, while right panel shows coloring by nCount. This Harmony-based batch correction and filtering approach was used to minimize donor-specific batch effects and remove cells with extreme expression.

**E.** RNA clustering (resolution = 0.5) showed less distinct clusters than ADT clustering.

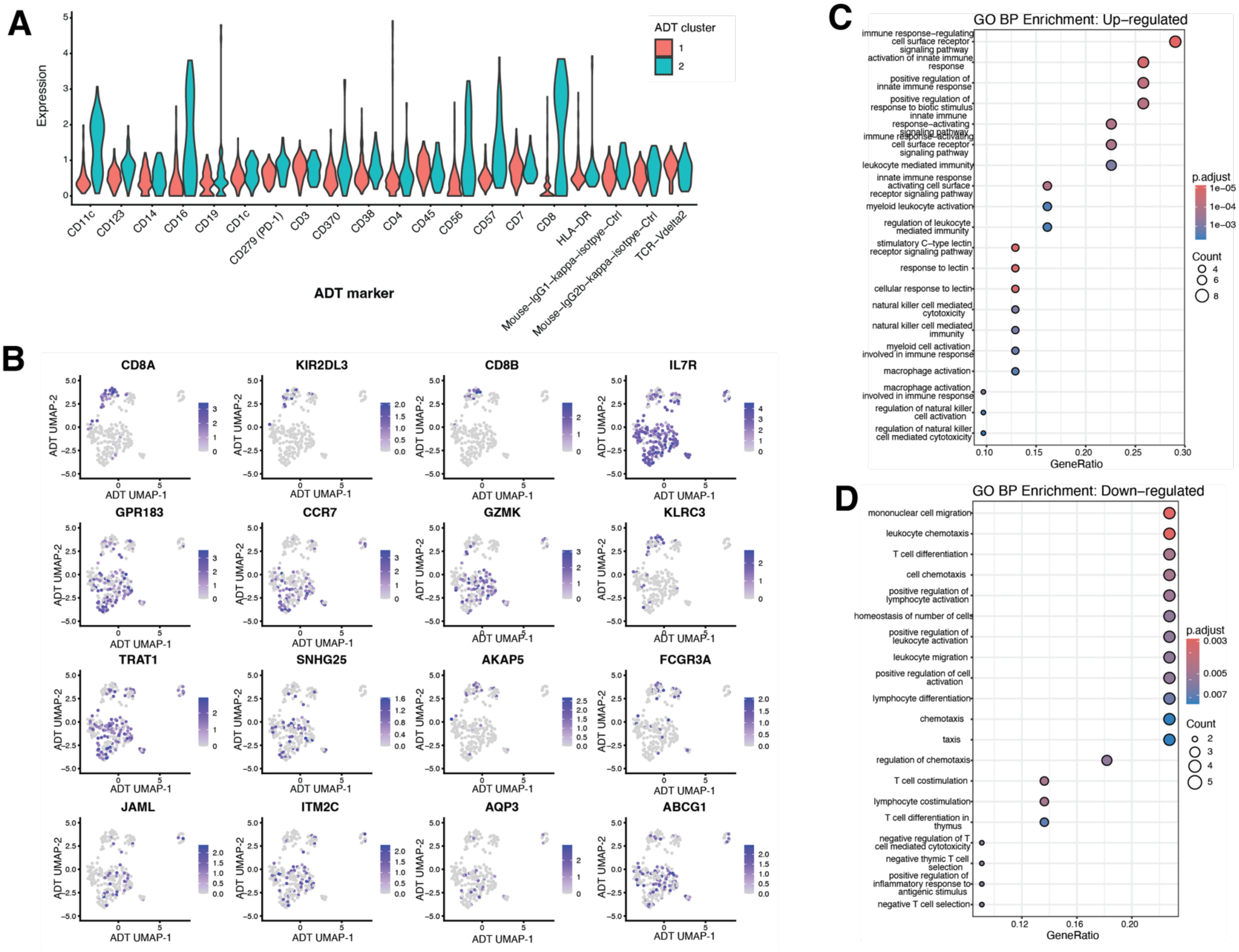

**Figure S9: CITE-seq differential gene expression by clusters**

**A.** Violin plot comparing ADT expression between 2 ADT clusters

**B.** Feature plots showing log1p normalized expression of representative differentially expressed genes across ADT-defined clusters

**C-D.** Gene Ontology (GO) biological process enrichment analysis of genes upregulated (C) or downregulated (D) in cluster 2 compared to cluster 1.

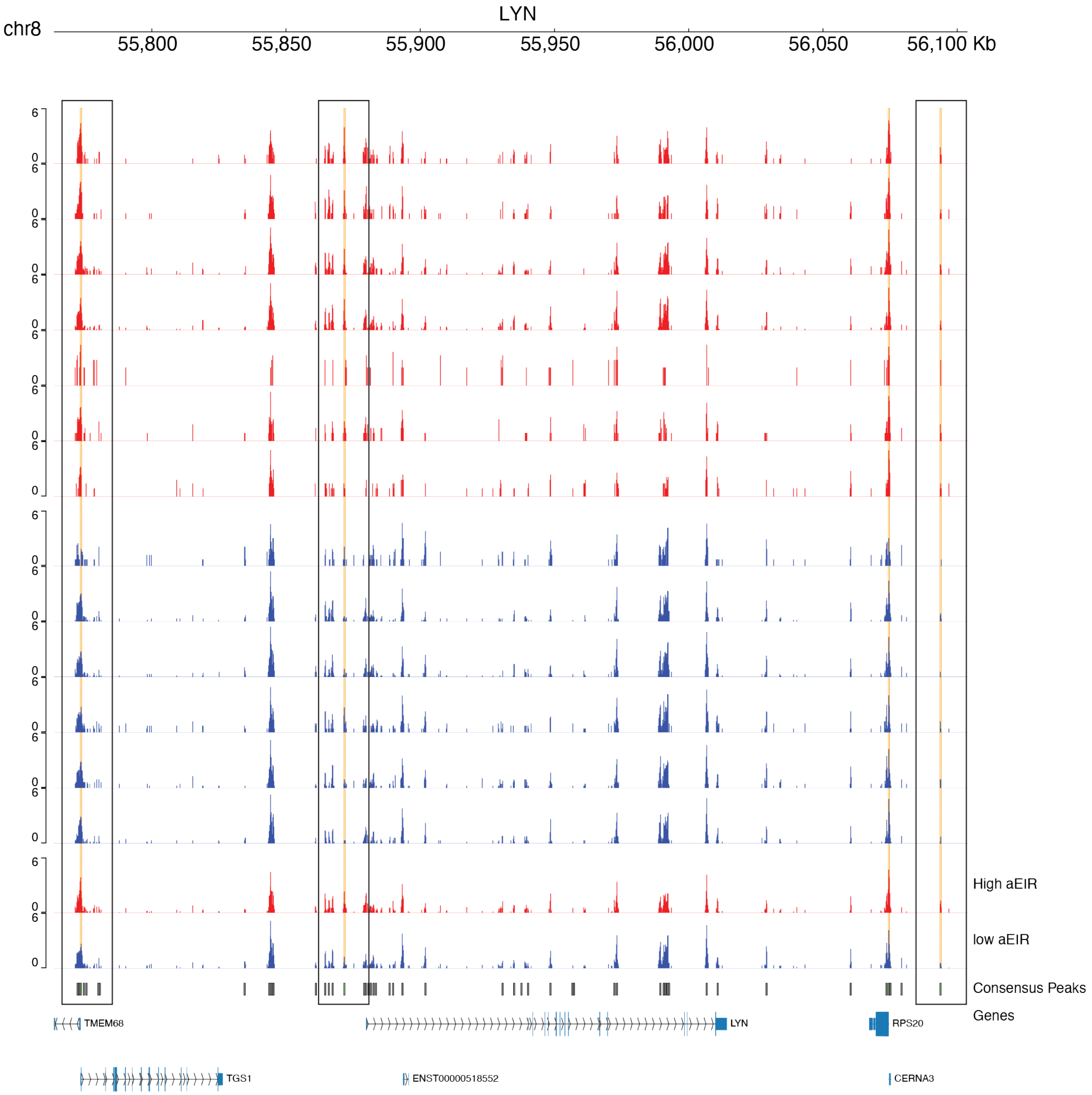


**Fig S10: Gene tracks for LYN across responder samples (N=13)**

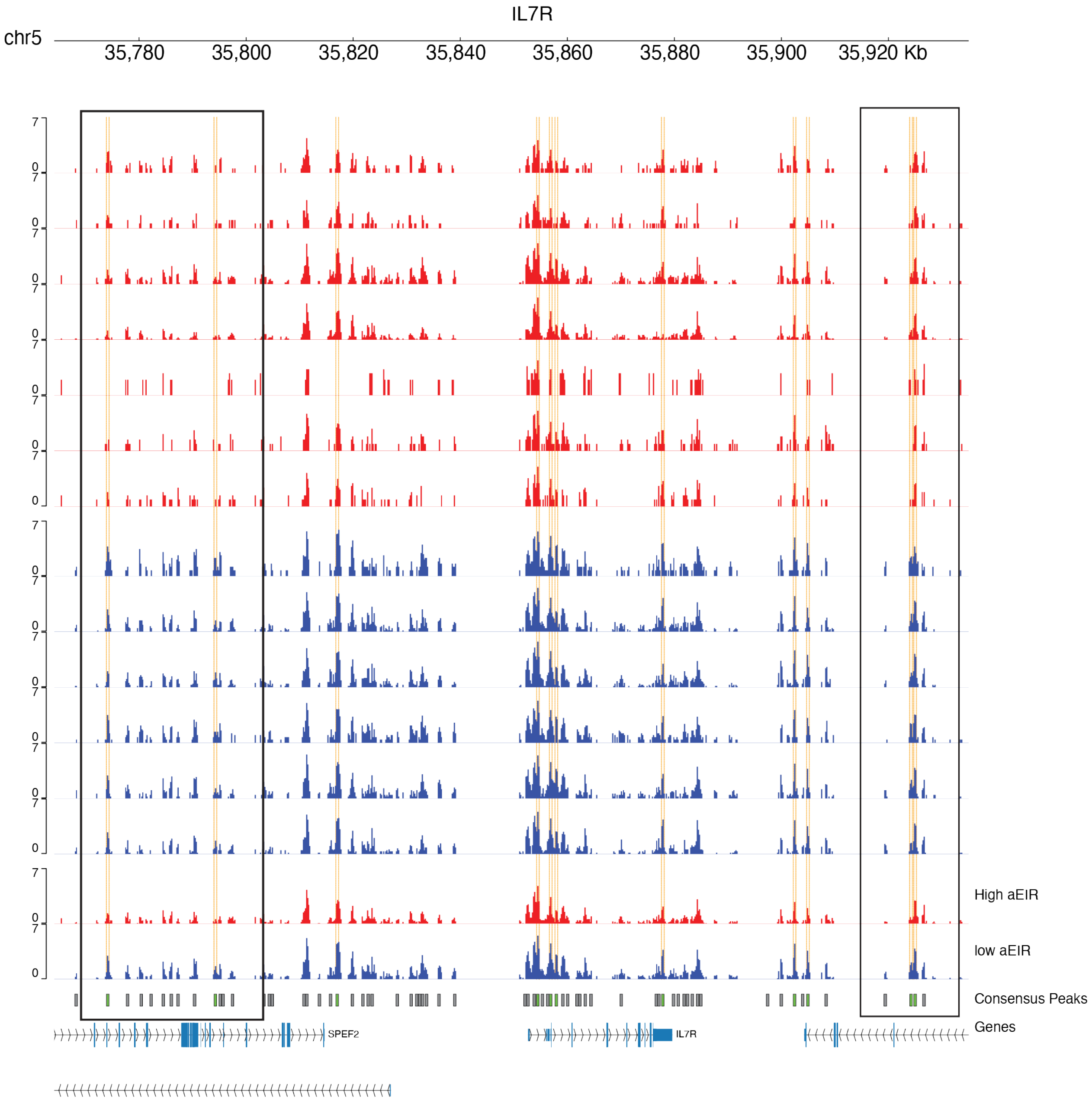


**Fig S11: Gene tracks for IL7R across responder samples (N=13)**

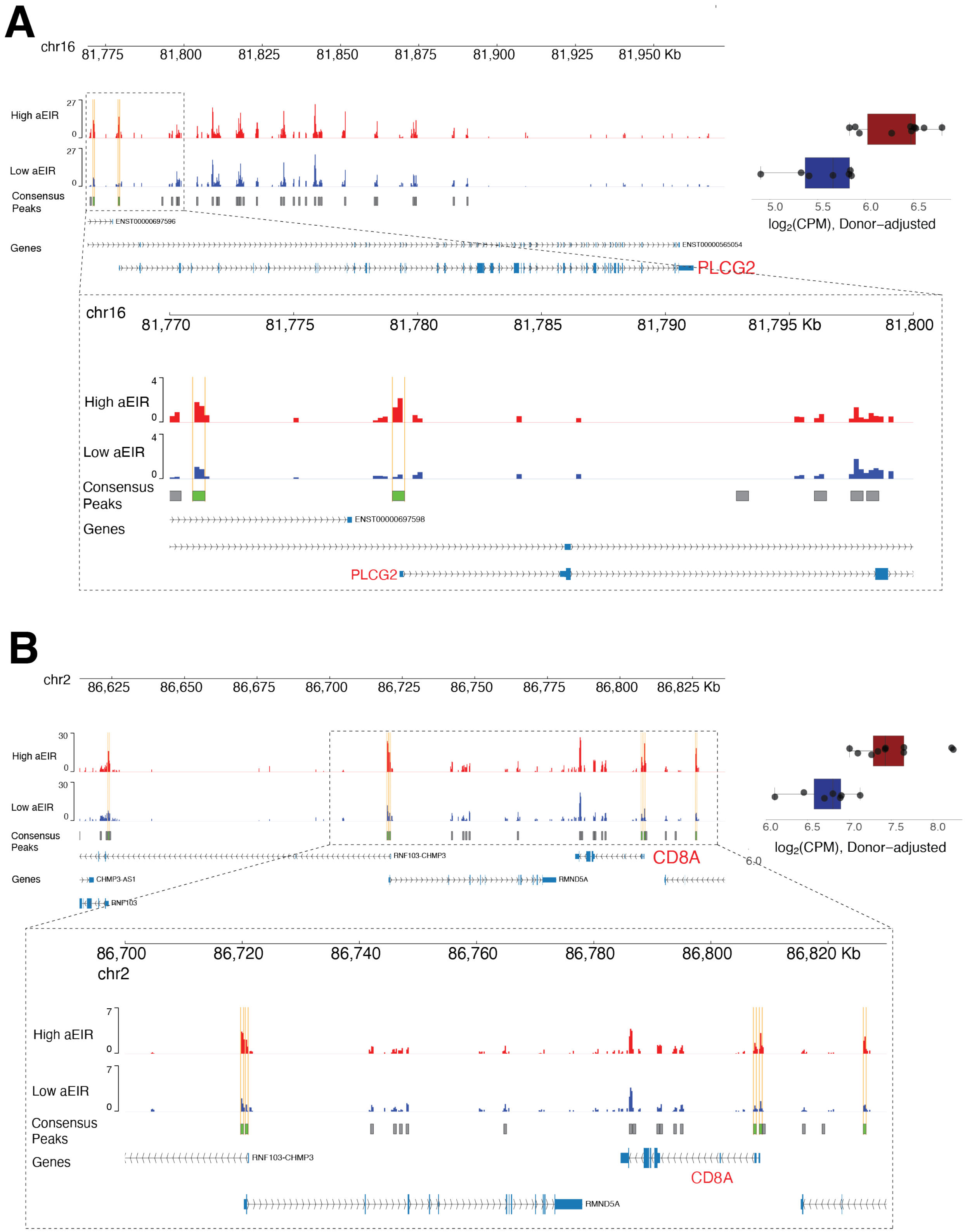


**Fig S12: Gene tracks for PLCG2 and CD8A**

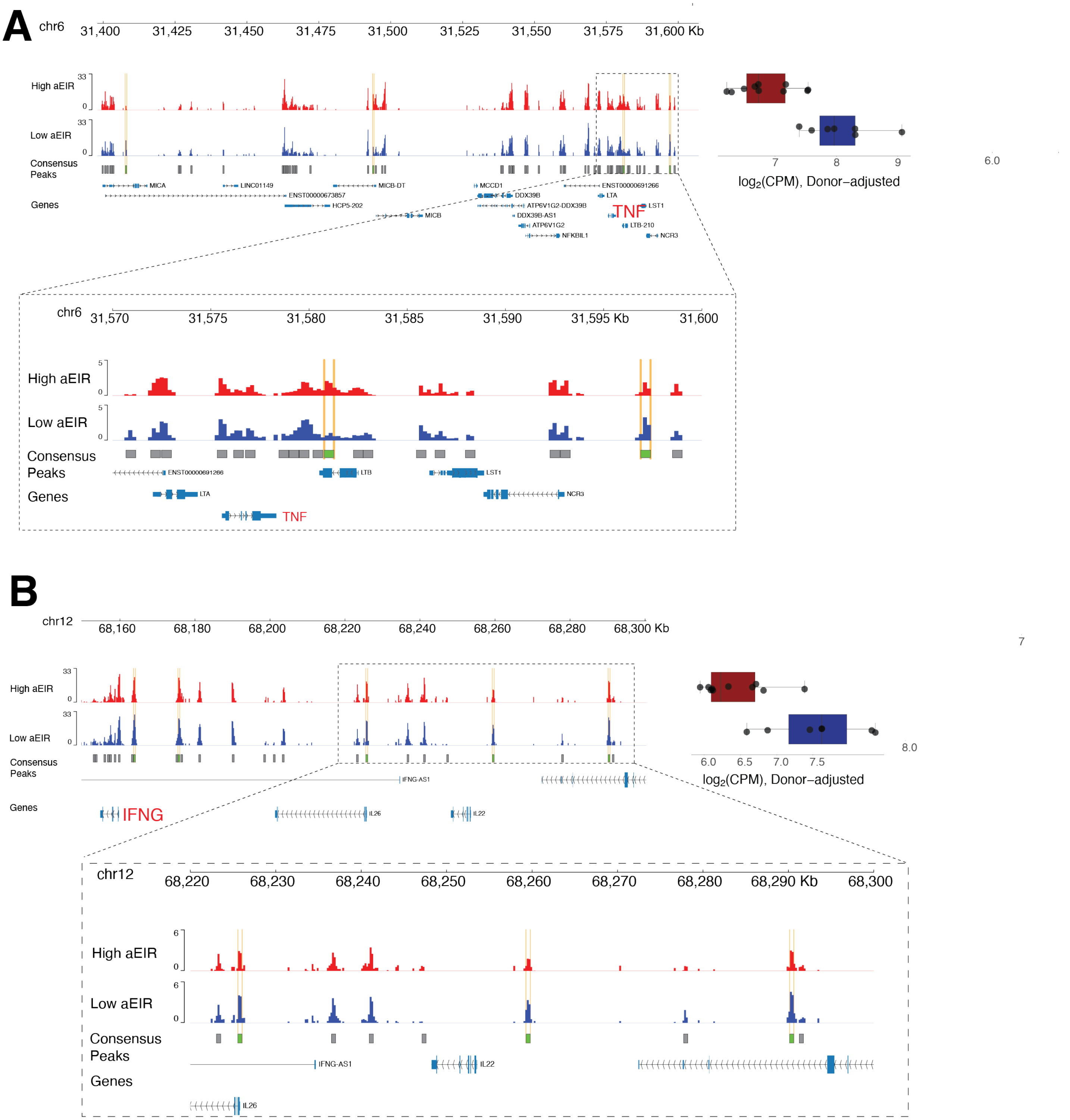


**Fig S13: Gene tracks for TNF and IFNG**

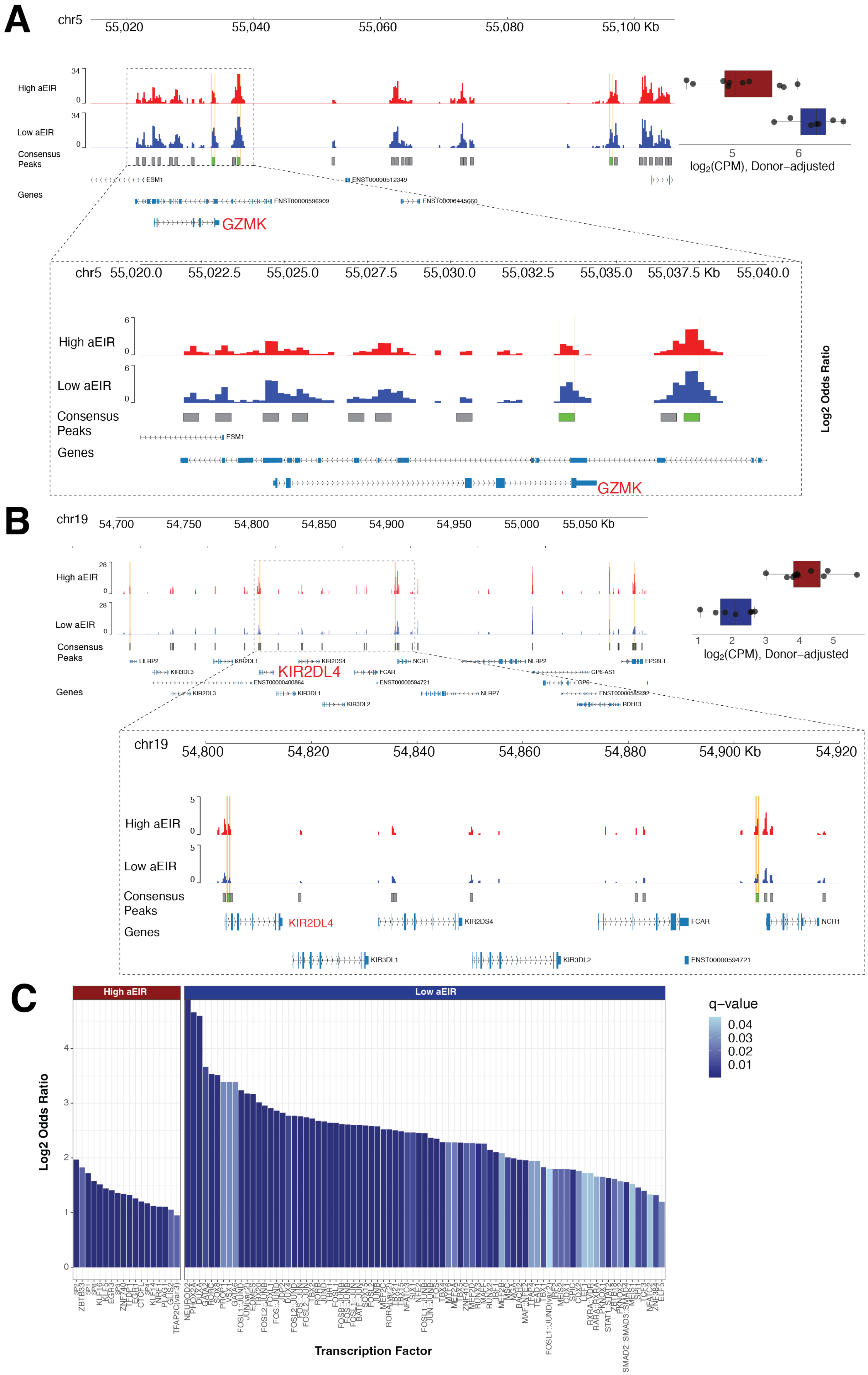

A
chr5
55,020
55,040
55,060
55,080
55,100 Kb
High aEIR
Low aEIR
Consensus Peaks
Genes
ESM1
ENST00000512349
ENST00000596909
ENST00000445660
GZMK
5
6
log2(CPM), Donor–adjusted
chr5
55,020.0
55,022.5
55,025.0
55,027.5
55,030.0
55,032.5
55,035.0
55,037.5 Kb
55,040.0
Log2 Odds Ratio
B
chr19
54,700
54,750
54,800
54,850
54,900
54,950
55,000
55,050 Kb
LILRP2
KIR3DL3
KIR2DL3
KIR2DL1
KIR2DL4
ENST00000400864
KIR3DL1
KIR2DS4
KIR3DL2
FCAR
ENST00000594721
NCR1
NLRP7
NLRP2
GP6-AS1
GP6
ENST00000585492
RDH13
EPS8L1
1
2
3
4
5
log2(CPM), Donor–adjusted
54,800
54,820
54,840
54,860
54,880
54,900 Kb
54,920
C
High aEIR
Low aEIR
q–value
0.04
0.03
0.02
0.01
Log2 Odds Ratio
Transcription Factor

**Fig S14: Additional gene tracks and LOLA enrichment for peak-to-gene linkage analysis**

**A.** Gene track for GZMK

**B.** Gene track for KIR2DL4

**C.** LOLA enrichment barplots of the linked peaks.

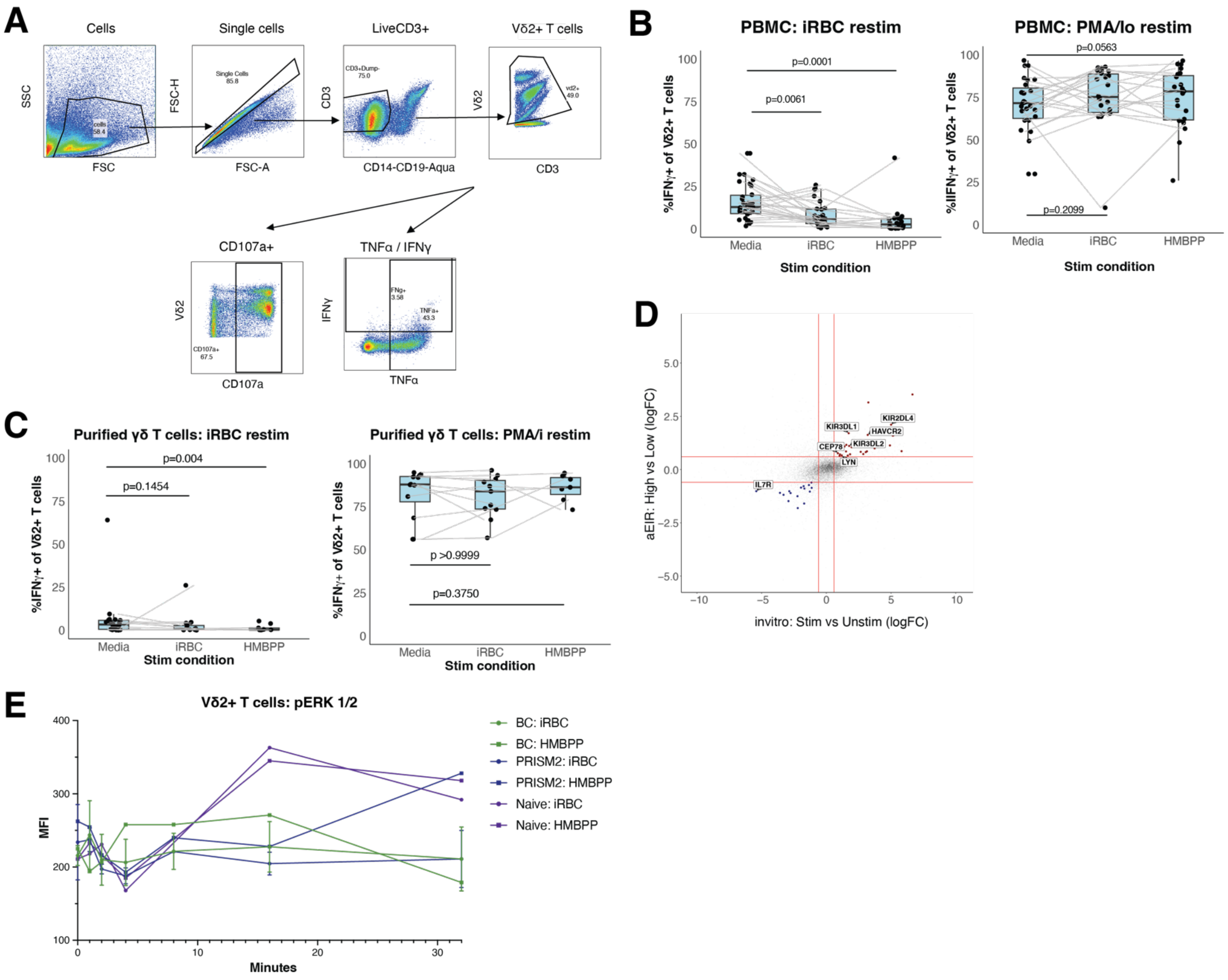


**Fig S15: Gating strategy and additional conditions for *in vitro* stimulation assays**

**A.** Gating strategy for Vδ2+ T cell *in vitro* stimulation assays showing representative PBMC sample.

**B-C.** Among Vδ2+ T cells within PBMCs (A) or purified Vδ2+ T cells (B), IFNγ expression decreases following 6-day iRBC or HMBPP stimulation followed by iRBC re-stimulation but do not change following re-stimulation with PMA/Ionomycin. Values normalized to media re-stim condition.

**D.** Overlap in RNA expression between longitudinal Ugandan samples and *in vitro*-stimulated Vδ2+ T cells.

**E.** Phosphoflow experiment showing Vδ2+ T cell pERK1/2 signaling in Uganda and naïve samples with diverse malaria exposure that were re-stimulated with iRBC.

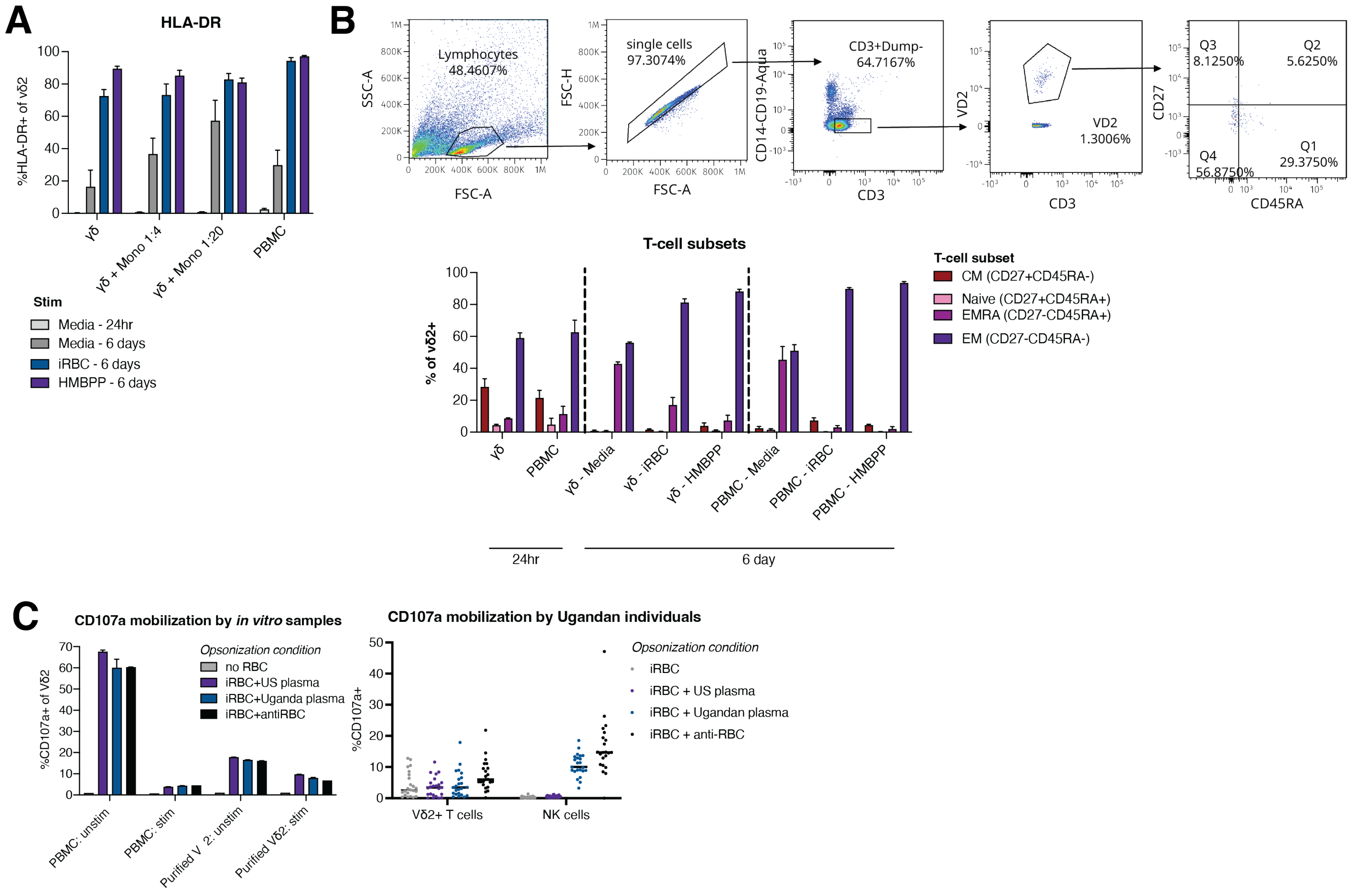


**Fig S16: Vδ2+ T cells responding to *in vitro* stimulation express HLA-DR and effector memory phenotypes and do not perform ADCC**

**A.** Expression of HLA-DR increases following stimulation with iRBC or HMBPP. N=3-4 depending on condition.

**B.** Gating strategy for defining memory subsets (top). Frequencies of effector memory (EM) cells increase following stimulation with iRBC or HMBPP, both for purified Vδ2+ T cells or Vδ2+ T cells within PBMCs (bottom). N=3-4 depending on condition.

**C.** Antibody-dependent cellular cytotoxicity assay using malaria-naïve samples (left) or Ugandan samples (right). iRBCs were opsonized with US or Ugandan plasma or controls and then added to PBMCs. Vδ2+ T cells do not show any evidence of ADCC while NK cells (right) do.

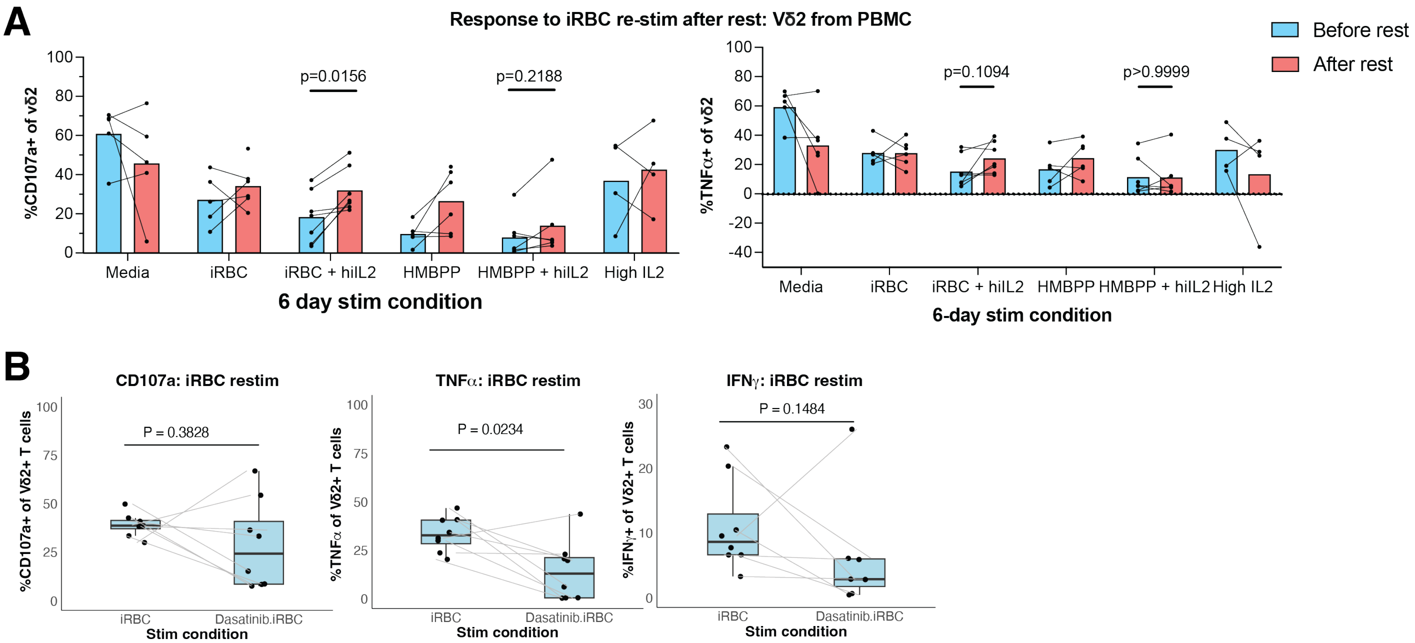


**Fig S17: Rest partially restores Vδ2+ T cell responses**

**A.** PBMCs were stimulated for 6 days with iRBC or HMBPP followed by a 3-5 day. Rest could partially rescue Vδ2+ T cell decreased responsivity to iRBCs. Values normalized to media re-stim condition.

**B.** Vδ2+ T cell CD107a, TNFα and IFNγ expression in PBMC samples with iRBC+Dasatinib compared to iRBC alone. Values normalized to media re-stim condition.
